# Copy number variation and elevated genetic diversity at immune trait loci in Atlantic and Pacific herring

**DOI:** 10.1101/2023.10.26.564141

**Authors:** Fahime Mohamadnejad Sangdehi, Minal S. Jamsandekar, Erik D. Enbody, Mats E. Pettersson, Leif Andersson

## Abstract

Atlantic and Pacific herring are sister species that diverged about 2 million years ago. Here we describe a genome comparison of the two species that reveal high genome-wide differentiation as expected for two distinct species but with islands of remarkably low genetic differentiation as measured by an *F_ST_* analysis. However, this is not caused by low interspecies sequence divergence but an exceptionally high estimated intraspecies nucleotide diversity. These high diversity regions are not enriched for repeats but are highly enriched for immune trait-related genes. This enrichment includes classical immunity genes, such as immunoglobulin, T-cell receptor and major histocompatibility complex genes, but also a substantial number of genes with a role in the innate immune system. An analysis of long-read based assemblies from two Atlantic herring individuals revealed extensive copy number variation at these genomic regions, indicating that the elevated intraspecies nucleotide diversities was partially due to the cross-mapping of short reads. This study demonstrates that copy number expansion and variation is a characteristic feature of immune trait loci in herring and that this genetic diversity is likely to contribute to resistance to infectious diseases in extremely abundant species, such as the Atlantic and Pacific herring. Another important implication is that these loci are blind spots in classical genome-wide screens for genetic differentiation using short-read data, not only in herring, likely also in other species harboring qualitatively similar variation at immune trait loci.

**Significance:** This study has revealed an extensive copy number variation and high nucleotide diversity at genes related to the immune system in Atlantic herring. Our analysis of previously published data in teleost species indicate that this is probably a widespread pattern among vertebrates. We also document that population genetic parameters estimated using short-read sequencing data are unreliable for these regions due to their complexity. They will also appear as blind spots in genome scans for regions of genetic differentiation based on *F_ST_* statistics due to the very high within population nucleotide diversity. We show how long-read data can be used to decipher the gene organization and genetic diversity at these regions.

## Introduction

Genetic screens based on whole genome sequencing are widely used to identify loci underlying variation in phenotypic traits and ecological adaptation. For instance, this approach has been successfully used to identify hundreds of loci in the Atlantic herring (*Clupea harengus*) with striking genetic differentiation associated with adaptation to different ecological conditions and timing of reproduction (Han et al. 2020). Genetic screens comparing different species may also be used to identify loci that explain phenotypic differences or have contributed to speciation, if the species are sufficiently closely related and do not show extensive differentiation across the entire genome. This approach has been used to identify loci involved in adaptation in adaptive radiations of closely related species of cichlids (McGee et al. 2020; Kautt et al. 2020) and Darwin’s finches (Lamichhaney et al. 2015; Rubin et al. 2022), among many others. Here we present a genomic comparison of the Atlantic herring and its sister species the Pacific herring (*Clupea pallasi*).

Atlantic and Pacific herring are both abundant species with key ecological roles in the North Atlantic and the North Pacific Oceans by being links between primary plankton production and carnivorous fish, sea birds and sea mammals. The Pacific herring is distributed along both the eastern and western sides of the Pacific Ocean, and in the Arctic Sea all the way until the White Sea (Semenova & Stroganov 2018). It spawns in shallow, near-shore habitats, in contrast with the deeper-spawning Atlantic herring (Hay et al. 2009). The two species separated about two million years ago (Martinez Barrio et al. 2016). However, there is a contact zone in the border between the Northeast Atlantic and the European side of the subarctic basin where gene flow occurs (Semenova & Stroganov 2018). Furthermore, a Pacific/Atlantic hybrid population is present in a subarctic fjord in Norway and has persisted for thousands of years (Pettersson et al. 2023).

Here, we performed a whole genome scan for genetic differentiation between Atlantic and Pacific herring to search for loci showing strong genetic differentiation between the two species. However, identification of loci that have contributed to evolutionary changes subsequent to speciation was hampered by high genome-wide sequence divergence. Surprisingly, the most striking discovery in this scan was the presence of loci that stood out due to remarkably low interspecific genetic differentiation, as indicated by *F_ST_* statistics. We show that these regions, referred to as high diversity regions, are highly enriched for genes involved in immune response and that the remarkably low *F_ST_* estimates are not due to low interspecies nucleotide divergence but by extremely high estimated intraspecies nucleotide diversity.

## Results

### Negative correlation between interspecies differentiation and intraspecies nucleotide diversity

We performed a genome-wide screening of genetic differentiation between Atlantic and Pacific herring, measured with *F_ST_*, *d_xy_* and *π* in 5 kb windows using short-read data from individually sequenced samples (Supplementary Table S1). First, average pairwise genetic divergence, estimated by *F_ST_*, was generally high between Atlantic and Pacific herring (genome average *F_ST_* = 0.58), most likely explained by accumulation of nucleotide substitutions and genetic drift subsequent to speciation and limited gene flow between the two species. This pattern is illustrated for chromosome 6 in Figure 1A (see Supplementary Figure S1 for all chromosomes). Second, *Fst* exhibits high variation across the chromosome, with most of the heterogeneity in *Fst* profile explained by variation in recombination rate; *Fst* values were negatively correlated (r = - 0.51, *P* < 0.001) with recombination rates (based on the Atlantic herring recombination map, Pettersson et al. 2019). Negative correlations between *Fst* and recombination rate have been reported in previous studies (Keinan and Reich 2010; Nachman and Payseur 2012; Wang et al. 2022). Stretches of the chromosome with inflated *F_ST_* values around the midpoint of the chromosome coincide with low recombining regions and we hypothesize that these are centromeric regions. Third, high *F_ST_* regions are interrupted by intervals of near-zero *F_ST_* levels, e.g., around 6.5 Mb on chromosome 6 (Figure 1A). Several shorter intervals with the same pattern are also present on this chromosome.

**Fig. 1.**
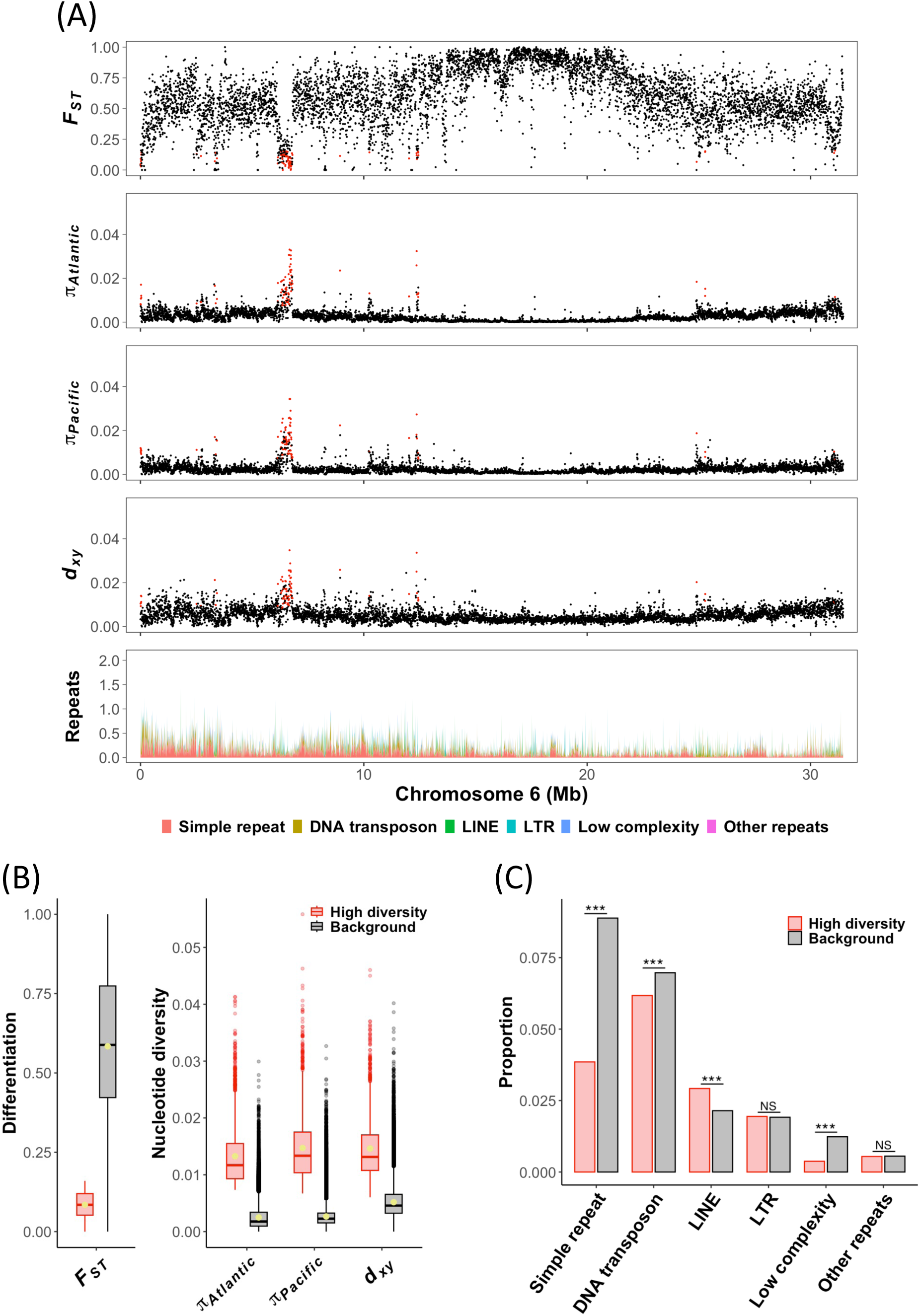
Distribution of population genetic parameters and repetitive elements calculated in non-overlapping 5 kb windows in a comparison of Atlantic and Pacific herring. **(A)** Estimates of population differentiation (*F_ST_*), intra-population nucleotide diversities in Atlantic (*π_Atlantic_*) and Pacific herring (*π_Pacific_*) and inter-population nucleotide diversity between Atlantic and Pacific herring (*d_xy_*) across chromosome 6 are displayed in the first to fourth tracks, respectively. Corresponding plots for all chromosomes are in Supplementary Figure S1. Each dot represents a 5 kb window. Windows in the top 5 percentile of genome-wide distribution of *F_ST_* and the bottom 5 percentile of genome-wide distribution of *π_Atlantic_* and *π_Pacific_* are highlighted in red. The bottom track represents the cumulative proportion of repeats across the chromosome. **(B)** The box plots show the distribution of genetic differentiation (*F_ST_*) and nucleotide diversities (*π* and *d_xy_*) in the high diversity regions (red) in comparison with genomic background (black); the mean values are represented with yellow dots inside the boxes. All comparisons are statistically significant (*P* < 0.001). **(C)** The bar chart summarizes the proportion of coverage by different repeat superclasses within the high diversity regions and background sequences. ***=*P* < 0.001; NS= non-significant.

Genome-wide averages of *π* estimated for the two populations were very close, with Pacific herring having a slightly higher level of nucleotide diversity (*π* = 0.0028) than Atlantic herring (*π* = 0.0026) (Figure 1A; Supplementary Figure S1). The average nucleotide diversity calculated in this study is comparable to previous reports (Fan et al. 2020; Atlantic *π* = 0.0030). The genome-wide average for pairwise nucleotide divergence (*d_xy_*) between Atlantic and Pacific herring was estimated at *d_xy_* = 0.0053 (Figure 1A and Supplementary Figure S1 show how this parameter varies across the genome). The U-shaped pattern of the *π* and *d_xy_* plots with low values in the middle of chromosome 6 most likely reflects variation in recombination rate; the correlation of recombination rate with *π_Atlantic_*, *π_Pacific_* and *d_xy_* was 0.45, 0.24 and 0.24, respectively (*P* < 0.001 for all comparisons). Further, we note that the regions with near-zero *F_ST_* are not associated with reduced interspecies sequence divergence but instead are driven by the very high denominator (within population *π*) that drastically reduces *F_ST_* estimates.

To further explore these signals and identify the mechanistic drivers of the observed pattern, we extracted windows with an *F_ST_* value lower than the 5th percentile (*F_ST_* < 0.16) and *π* values above the 95th percentile in each species (*π_Atlantic_* > 0.0073 and *π_Pacific_* > 0.0067). Based on these criteria, 2,030 windows (∼10 Mb in total) with particularly low *F_ST_* and high nucleotide diversity compared with genomic background were identified (Figure 1B). Throughout the paper, these regions are designated as high diversity regions. *F_ST_* has a moderate negative correlation with intraspecies nucleotide diversities estimated for the whole genome (−0.56 with *π_Atlantic_* and −0.42 with *π_Pacific_*; *P* < 0.001) but this relationship weakens in the high diversity regions (−0.21 and −0.24, respectively; *P* < 0.001) (Supplementary Figure S2). Per-window estimates of nucleotide diversity within Atlantic and Pacific herring show a strong correlation (0.74; *P* < 0.001), as expected for closely related species. *d_xy_* exhibits similar patterns to *π*, as seen in the distribution plot, and a strong positive correlation exists between *d_xy_* and *π* in each species (0.77 with *π_Atlantic_* and 0.70 with *π_Pacific_*; *P* < 0.001); this correlation is even stronger in the high diversity regions (0.91 and 0.92, respectively; *P* < 0.001).

### Repetitive elements are not enriched in high diversity regions

Minimal interspecies differentiation at the high diversity regions implies that a large proportion of the total variation occurs within species and thus, these signals can potentially be footprints of balancing selection. Balancing selection is characterized by maintaining genetic diversity above neutral expectations, thereby leading to low levels of genetic differentiation between populations at regions under selection (Fischer et al. 2014; Brandt et al. 2018). However, an alternative hypothesis is that the high diversity regions are enriched for repetitive elements and the detected signals represent an artefact of the mapping of short sequence reads. Due to high copy number and sequence identity inherent in repetitive elements, these difficult-to-assemble regions of the genome tend to yield unreliable SNP calls. To test if the signals were due to interference from repeats, we calculated the proportion of each window occupied by different classes of repetitive elements. This analysis showed that the high diversity regions are not enriched for repetitive elements (Figure 1A). On the contrary, the average proportion of total repeats observed at the high diversity regions (0.16) is lower than the genomic background (0.22; Welch’s t-test, *P* < 0.001). More specifically, simple repeats and low complexity repeats are less frequent across the high diversity regions, compared with the rest of the genome (0.039 and 0.004 vs. 0.089 and 0.012, respectively; Welch’s t-test, *P* < 0.001) (Figure 1C). However, we noted that many of the high diversity regions in our study are flanked by regions with a high density of repetitive elements, particularly simple sequence repeats in their immediate surroundings, although the frequency of these elements drops inside the regions themselves.

### High diversity regions are highly enriched for immune-related genes

We performed gene ontology (GO) term overrepresentation analysis in order to characterize the genes located in the high diversity regions. Out of 24,095 protein-coding genes in the herring genome, 881 genes had full or partial overlap with the high diversity regions. Only the genes annotated with at least one GO term were incorporated in the analysis, leading to a total of 21,345 genes retained, out of which 587 overlap the high diversity regions; 17,783 genes genome-wide and 270 genes in the high diversity regions had one or more biological process (BP) term annotations assigned. Thus, genes located in the high diversity regions are more often unannotated than the genome average (0.69% vs 0.26% for BP terms; χ^2^ test, *P* < 0.001). Keeping only GO terms associated with a minimum of ten genes, 6,690 BP terms were assessed. The overrepresentation analysis was conducted with Weight01 algorithm implemented in the topGO R package (Alexa & Rahnenfuhrer 2010), and revealed that the high diversity regions are highly enriched for genes involved in immune-related processes (Figure 2). This algorithm accounts for GO dependencies and hierarchical structure (Alexa et al. 2006), but there are nevertheless some degrees of overlap between top GO terms. These overlapping GO terms are closely related in the GO hierarchy. The interrelationships and connections among significant terms are illustrated in a GO graph in Supplementary Figure S3A. The genes labelled with each term, and summary statistics of the GO terms, are compiled in Supplementary Table S2. Overrepresentation results for molecular function (MF) and cellular component (CC) terms are presented in Supplementary Table S2 and Supplementary Figure S3 (B and C).

**Fig. 2.**
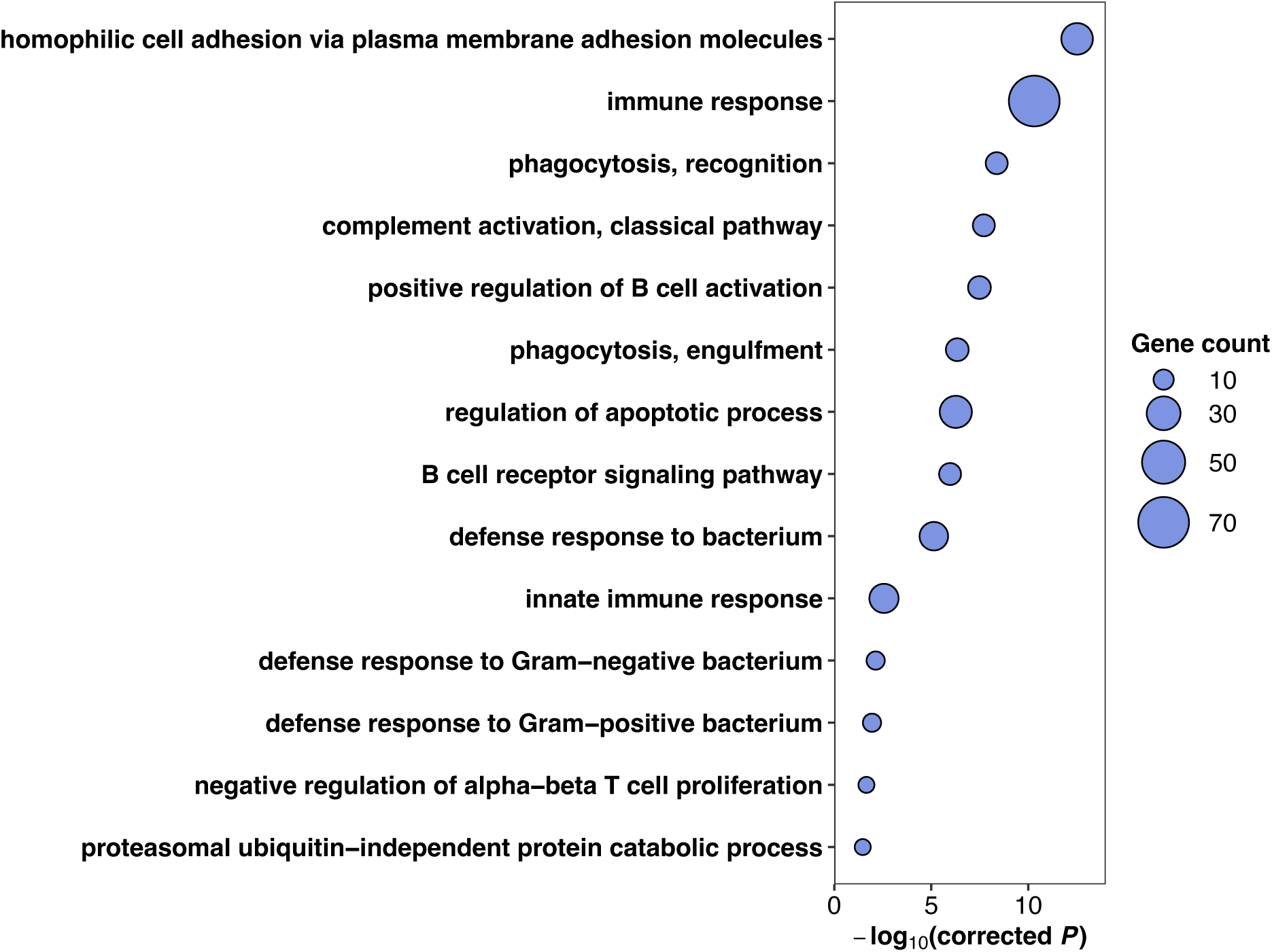
Significant biological process terms from GO overrepresentation analysis (*P* ≤ 0.01; corrected for multiple testing with the Bonferroni method) for genes present in high diversity regions in herring. The size of each circle is proportional to the number of enriched genes in the corresponding GO category.

The above findings encouraged us to conduct a more detailed examination of some of the high diversity loci to explore structure and organization of the genes. For example, a 400 kb region around position 6.5 Mb on chromosome 6 comprises a cluster of genes coding for proteins that contain Immunoglobulin-like domains (Figure 3). These genes are homologous to Novel immune-type receptor (*NITR*) genes in other teleosts (Yoder et al. 2001; Desai et al. 2008). Additional examples of gene organization in high diversity regions showing a similar pattern are presented in Supplementary Figure S4 and a comprehensive list of high diversity loci can be found in Supplementary Table S3. Based on these analyses we conclude that the most characteristic feature of the high diversity regions in the comparison of Atlantic and Pacific herring is that they are composed of clusters of immune-related genes.

**Fig. 3.**
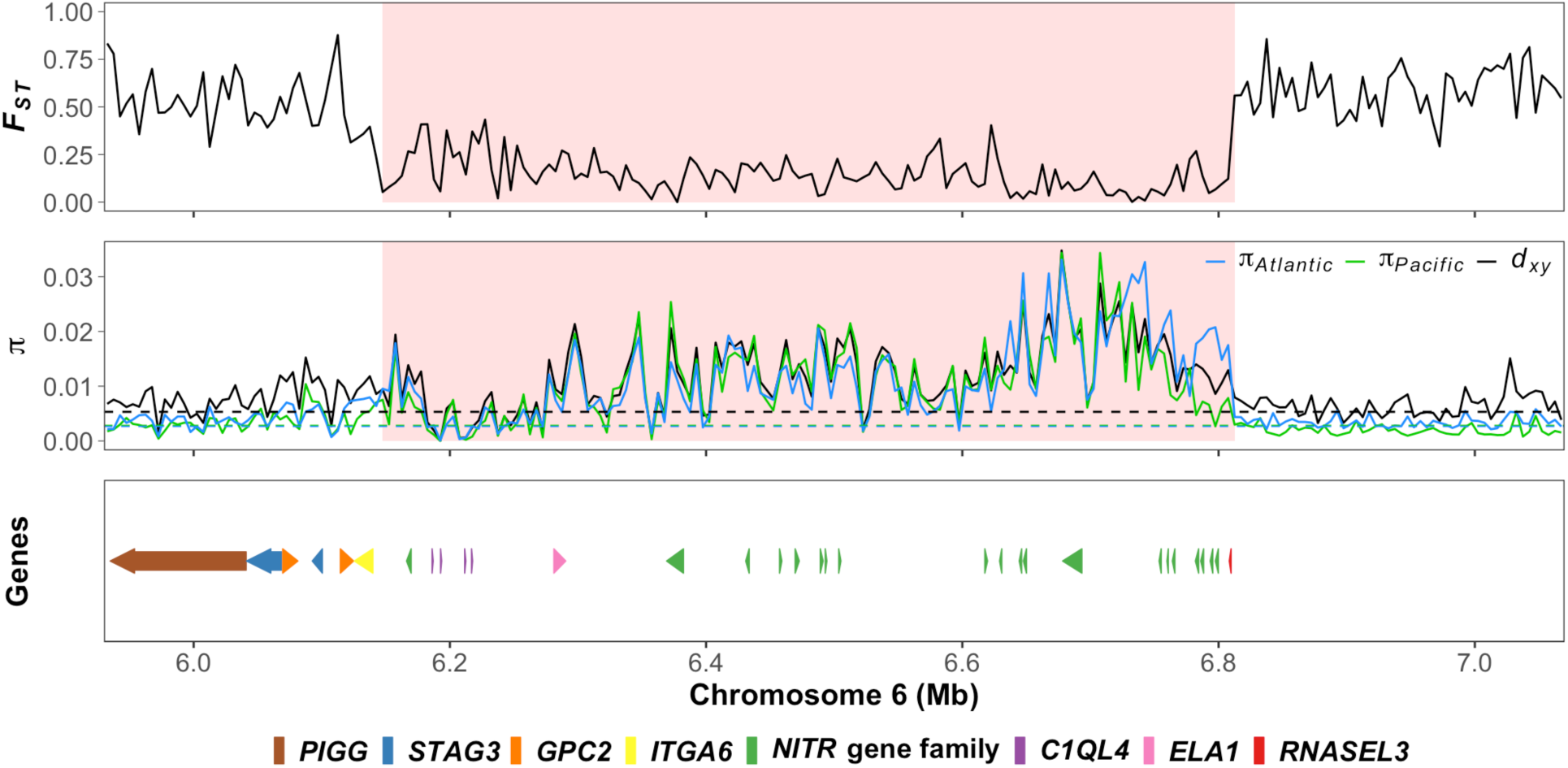
Zoom-in plots of a high diversity region on chromosome 6. The first track shows a marked *F_ST_* drop in the highlighted region. The second track shows the elevated *π_Atlantic_*, *π_Pacific_* and *d_xy_* in the same region; dashed lines depict genome-wide averages. The shaded area in the plots depicts merged high diversity regions. Gene organization on the reference assembly is presented in the bottom track (annotation source: NCBI Clupea harengus Annotation Release 102), indicating a cluster of *NITR* genes within the high diversity region. Color code for genes is given below the figure.

To assess the prevalence of the observed pattern in natural populations of other species, we looked for signals of low differentiation and high nucleotide diversity in previously published data from fish. We extensively searched for genome scan studies focusing on adaptive radiations occurring during later stages of speciation. Among the limited number of such studies, we identified comparable signals among radiations of Midas cichlid (Kautt et al. 2020) and stickleback (Yamasaki et al. 2020) that contain clusters of immune trait genes similar to the ones observed in herring (Supplementary Table S4), suggesting that despite being a repeated pattern in teleost fish, this has been largely overlooked, or at least left undescribed, in previous comparative genomic studies.

### PacBio long-read data reveal extensive copy number variation at high diversity regions

The analysis of genetic diversity at clusters of closely related genes using short-read data is challenging, partially because the genome assembly may be incorrect with collapsed copy number variation and partially because of the difficulty to align short reads to the correct copy – in fact, due to structural variation, there is often no “correct copy” for a subset of reads. We therefore generated *de novo* genome assemblies of two Atlantic herring individuals based on PacBio HiFi long reads (Table 1) to overcome this problem. This allowed us to study copy number variation as well as to more accurately estimate levels of nucleotide diversity. Here we present a detailed analysis of three representative genomic regions with low *F_ST_* containing clusters of three gene families: immunoglobulin heavy variable (*IGHV*), CD300e molecule (*CLM2*), and interferon-induced protein with tetratricopeptide repeats 10 (*IFIT10*). We first annotated the genes and noted the presence of multiple copies along with pseudogenes composed of incomplete or disrupted coding sequences (Table 2). The *IGHV* locus varied between 30 and 70 gene copies distributed across a region ranging in estimated size from 64 kb to 160 kb on different haplotypes (Figure 4A). Copy number variation was also evident from the fact that coverage values were elevated relative to the average coverage across the genome (Figure 4B).

**Table 1.**
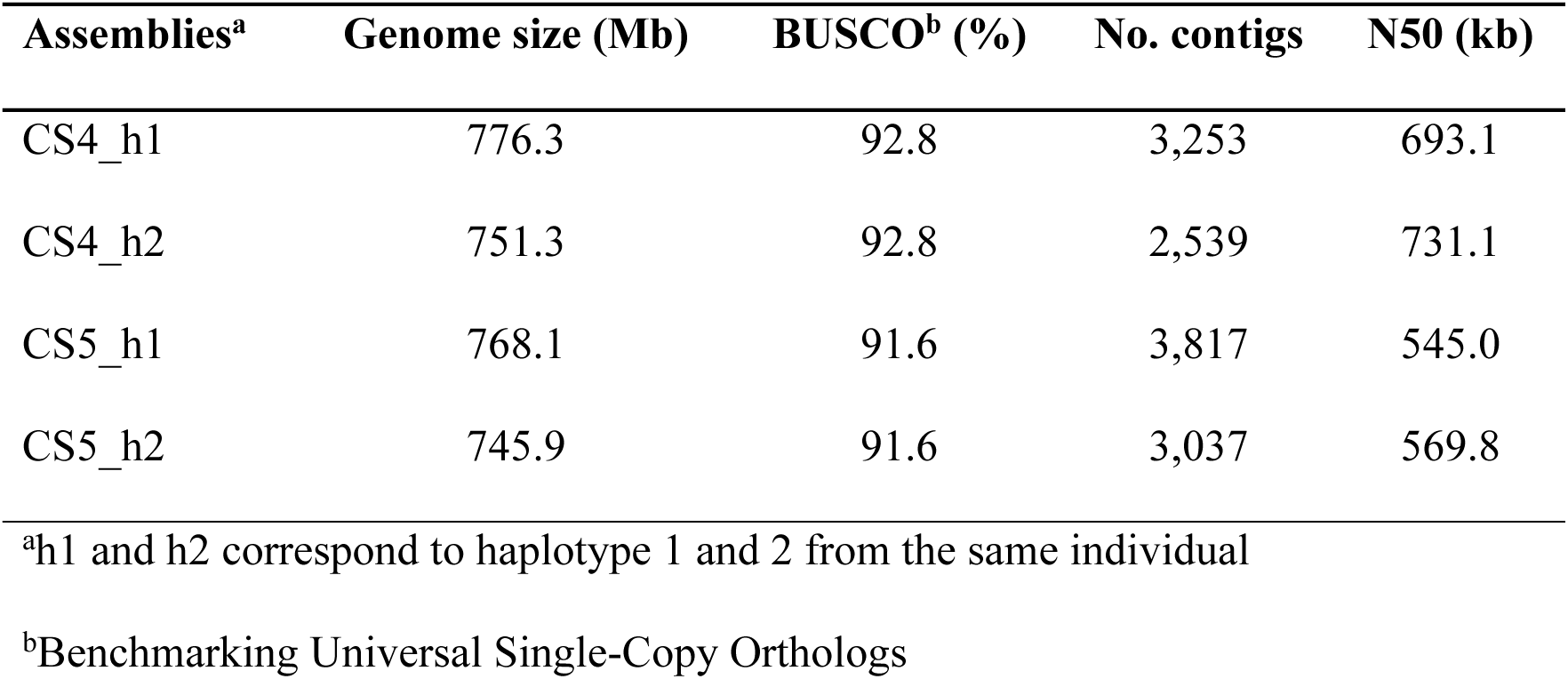
Statistics for haplotype-phased genome assemblies of two Atlantic herring individuals (CS4 and CS5) based on PacBio HiFi sequencing.

**Table 2.** Comparison of the number of gene copies of *IGHV*, *CLM2*, and *IFIT10* in three high diversity regions among five assemblies, the reference assembly (Pettersson et al. 2019) and four PacBio haplotype assemblies (present study). The total number of gene copies in each region is indicated as functional copies plus the number of obvious pseudogenes (**ψ**) given in parentheses, the latter carry inactivating mutations.

| Assemblies | *IGHV* |  | *CLM2* |  | *IFIT10* |  |
| --- | --- | --- | --- | --- | --- | --- |
| | No. of<br>genes ($\psi^a$) | size (kb) | No. of<br>genes ($\psi^a$) | size (kb) | No. of<br>genes ($\psi^a$) | size (kb) |
| Reference | 40 (0) | 94 | 4 (1) | 29 | 2 (0) | 22 |
| CS4_h1 | 30 (0) | 70 | 4 (3) | 47 | 2 (1) | 42 |
| CS4_h2 | 70 (0) | 160 | 4 (1) | 53 | 2 (1) | 49 |
| CS5_h1 | 40 (0) | 100 | 4 (4) | 74 | 2 (2) | 58 |
| CS5_h2 | 34 (0) | 64 | 4 (4) | 66 | 2 (1) | 29 |

<sup>a</sup> $\psi$ = Number of pseudogenes

**Fig. 4.**
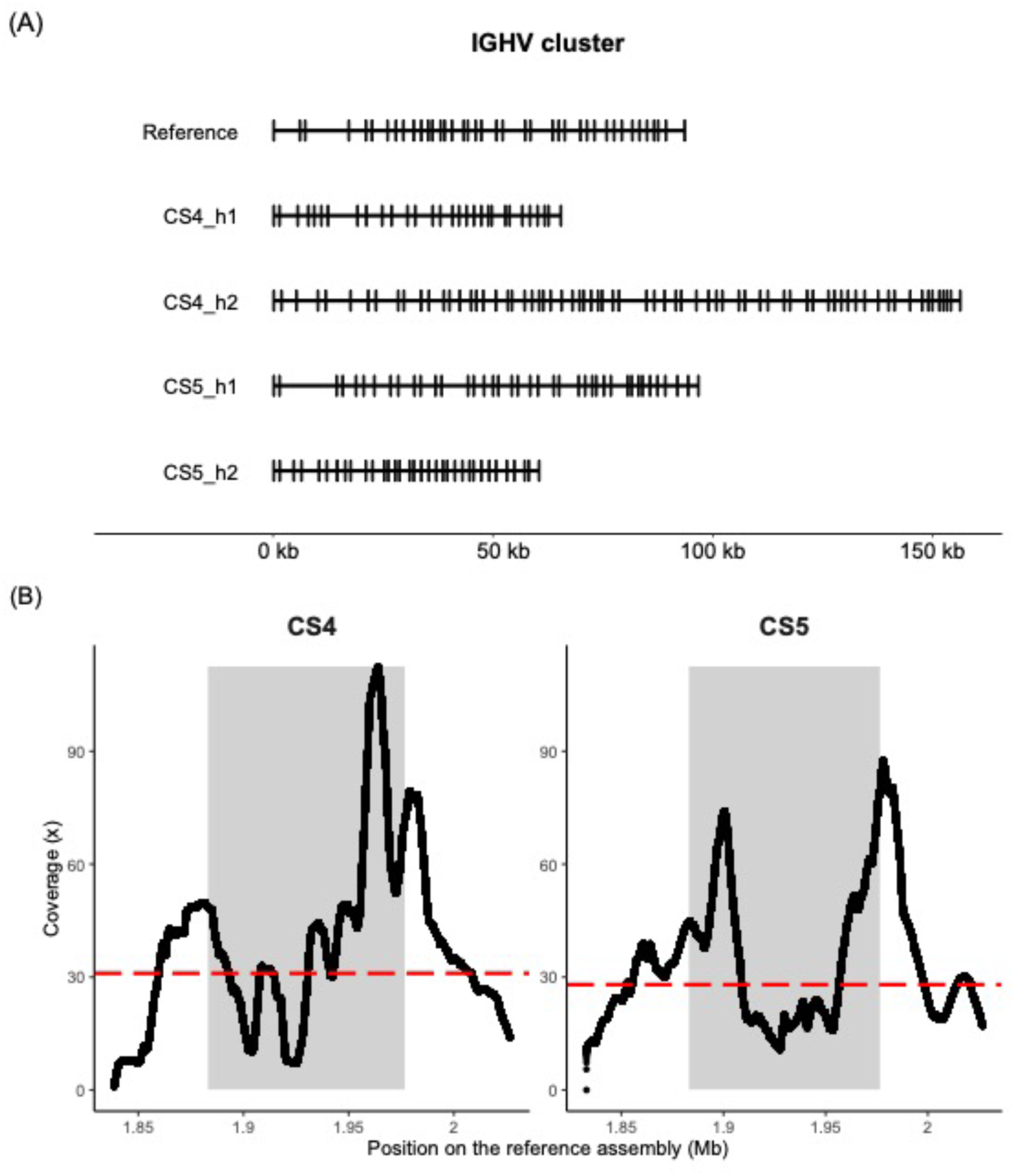
Copy number variation of the *IGHV* genes on chromosome 1 in the region 1,882,931-1,976,669 bp in the reference assembly. **(A)** Genome organization of the *IGHV* region on the Atlantic herring reference assembly and on four PacBio-based haplotype assemblies. **(B)** Sequence coverage obtained after aligning PacBio HiFi reads from two individuals (CS4 and CS5) to the reference herring genome assembly. The dots represent rolling average coverage in 10 kb window. Shaded area corresponds to the *IGHV* region. Average genome coverage is indicated by a dashed red line.

The Atlantic herring reference assembly (Pettersson et al. 2019) contains a cluster of four *CLM2* genes and one *CLM2* pseudogene on chromosome 22. The PacBio assemblies of this region also contained four full length copies and the number of CLM2 pseudogenes varied from one to four (Figure 5). Furthermore, the length of the PacBio assemblies were all longer than the reference assembly suggesting that duplicated sequences may have been collapsed in the reference assembly. A phylogenetic tree analysis revealed that the *CLM2C* sequences formed a distinct group whereas the genes designated *CLM2A*, *CLM2B1*, and *CLM2B2* were all closely related and did not form three distinct allelic series. Thus, it is not possible to align short read sequences to the correct copy (based on genomic location) of *CLM2A/B* genes. The assembled region containing a pair of *IFIT10* genes on chromosome 23 were all longer on the new PacBio assemblies compared with the reference assembly and the former contained an *IFIT10* pseudogene lacking in the reference assembly (Figure 6).

**Fig. 5.**
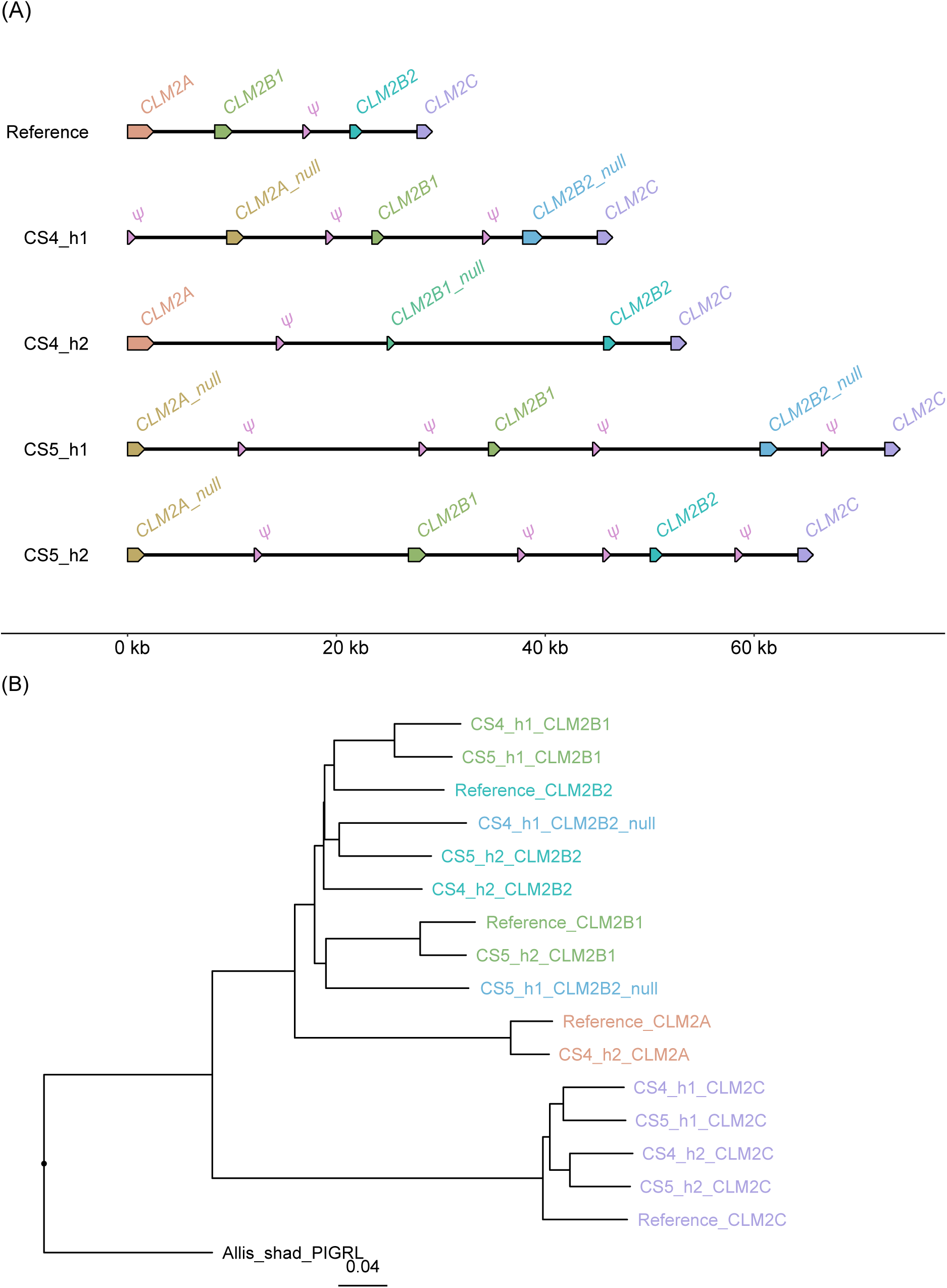
Copy number variation of the cluster of *CLM2* genes on chromosome 22 in the region 18,433,425–18,434,439 bp of the reference assembly. **(A)** Genomic organization of *CLM2* genes on the Atlantic herring reference assembly and four PacBio-based haplotype assemblies. *CLM2A_null* lacks exon1 and intron1 which is present in *CLM2A*. Similarly, *CLM2B1_null* in CS4_h2 lacks exon1 and intron1 which is present in *CLM2B1*. *CLM2B2_null* in CS4_h1 and CS5_h1 has one nonsense mutation in exon2. Sequences referred to as pseudogenes (*ψ*) contain only fragments of *CLM2* coding sequences. **(B)** Phylogenetic tree of *CLM2* coding nucleotide sequences. The color code is the same as in Figure 5A.

**Fig. 6.**
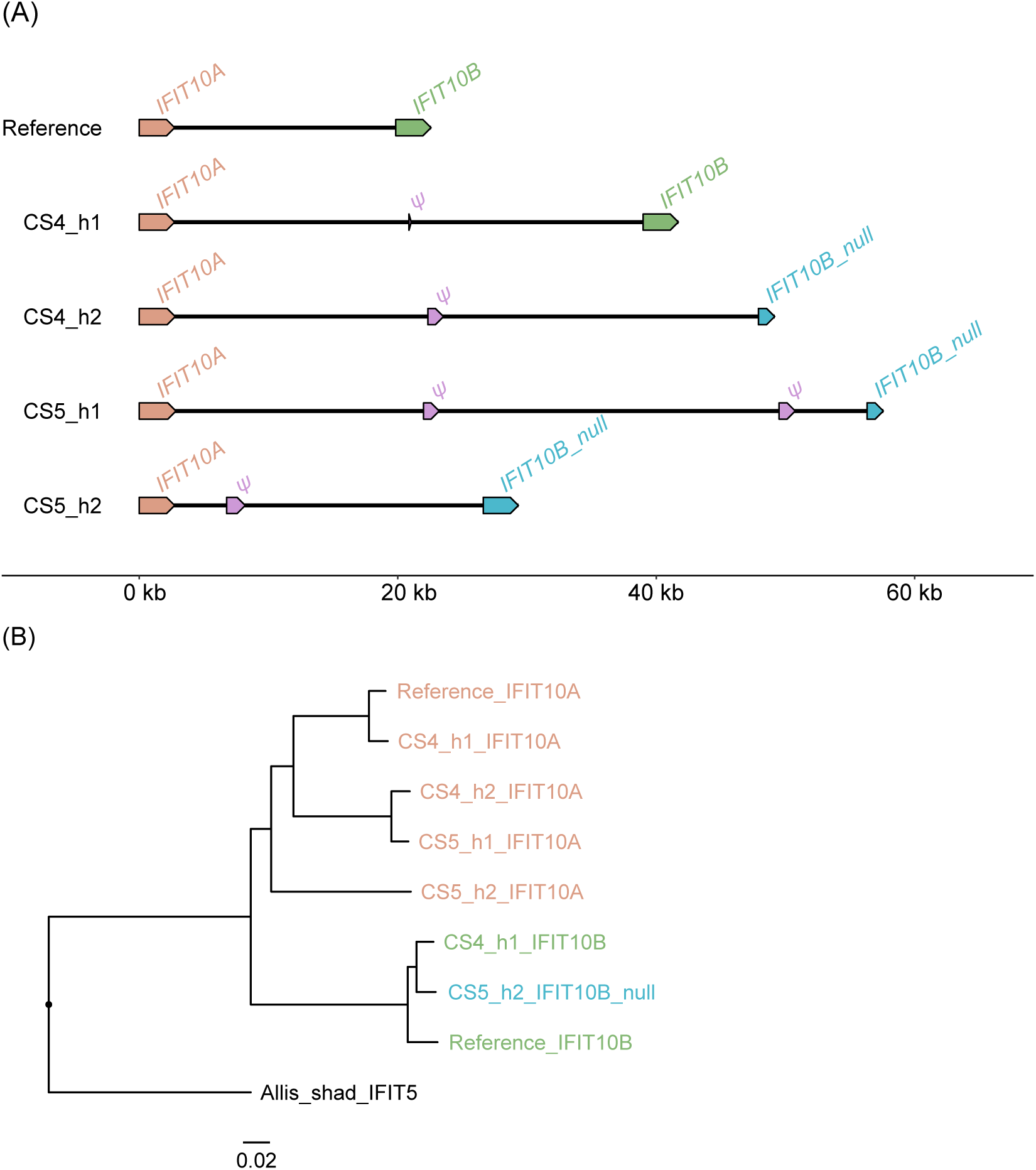
Copy number variation of the cluster of *IFIT10* genes on chromosome 23 in the region 12,114,407-12,116,709 bp of the reference assembly. **(A)** Genomic organization of *IFIT10* genes on five haplotypes. *IFIT10B*_*null* in CS4_h2 and CS5_h1 had multiple stop codons throughout the gene length. *IFIT10B_null* has a deletion of four bp. **(B)** Phylogenetic tree of *IFIT10* coding nucleotide sequences with Allis shad *IFIT5* as an outgroup. The color code is the same as in Figure 6A.

We estimated nucleotide diversity (*π*) only for the coding sequences to ensure that allelic sequences were compared. This revealed considerably lower *π* values than estimated by short-read sequencing data, although several of these genes had still much higher nucleotide diversity than the genome average of 0.003 (Supplementary Table S5 for *IGHV* and Supplementary Table S6 for *CLM2* and *IFIT10*).

## Discussion

In the current study, a window-based scan of sequence divergence between Atlantic and Pacific herring revealed an unexpected set of low-*F_ST_*, high-*d_xy_* regions that clearly stand out against the genomic background, which is characterized by high *F_ST_* and moderate *d_xy_*. Close inspection of these high diversity regions unveiled a striking enrichment of immune-related genes, typically forming multigene family clusters. To our knowledge, no previous study has primarily focused on such regions, but by re-examination of publicly available data, we found similar signals among Midas cichlids (Kautt et al. 2020) and sticklebacks (Yamasaki et al. 2020). These findings imply that the observed pattern is widespread, yet has remained largely unexplored as regards its contribution to genome evolution during speciation and adaptation.

We show that clusters of genes involved in both adaptive and innate immunity are highly enriched at the high diversity regions detected in this study. Immune response genes are among the fastest evolving genes and species/lineage-specific clades of different categories of immune genes are commonly observed (Hill et al. 2019). This is a reasonable explanation as to why a relatively large proportion of the genes in the high diversity regions lack functional annotation. Furthermore, the presence of several pseudogenes in the multigene clusters is consistent with these loci following a gene birth-and-death model (Nei et al. 1997; Nei and Rooney 2005). Many of the genes present in the high diversity regions belong to the innate immune system, an observation consistent with the fact that in fish, defense mechanisms against pathogens are skewed towards innate immunity (Magnadóttir 2006; Uribe et al. 2011), while in mammals, adaptive immune responses are more important for immune protection (Litman et al. 2010; Han et al. 2020b). Previous studies have also noted lineage/species-specific gene family expansions of genes involved in innate immunity in teleosts relative to other vertebrates, and related these expansions to the evolutionary success of the teleost lineage; this includes genes for complement factors (Najafpour et al. 2020), toll-like receptors (*TLR*) (Star et al. 2011; Solbakken et al. 2016), novel immune-type receptor (*NITR*) (Desai et al. 2008; Ferraresso et al. 2009), NACHT-domain and leucine-rich-repeat-containing (*NLR*) (Howe et al. 2016; Suurväli et al. 2022), tripartite motif proteins (*TRIM*) (Boudinot et al. 2011; Suurväli et al. 2022), CC chemokines (Liu, Tingyu Wang, et al. 2020) and interferons (Liu, Tiehui Wang, et al. 2020). Our observations demonstrate that immune-related gene families also have experienced rapid and dynamic evolution in the two sister species of herring studied here.

Studying genomic regions harboring multigene family clusters proves to be challenging with the use of short sequence reads because of the difficulty to distinguish allelic sequences from paralogs. Therefore, population genetic parameters, such as *F_ST_*, estimated based on short-read sequencing data are prone to be biased at such regions, and hence should be interpreted with caution. We therefore used PacBio long-read data to make it possible to separate haplotypes of homologous chromosomes and identified fine-scale pattern of these complex regions. This analysis revealed copy number variation (CNV), and other re-arrangements, at many of the high diversity loci. In fact, none of the four haplotypes deduced from two individuals had an identical structure to another haplotype at the three gene regions studied in detail (Figures 4-6). The analysis showed that the extensive genetic diversity at immune-related genes in the Atlantic herring is due to a combination of copy number variation and nucleotide diversity between alleles.

Previous studies have detected a large number of loci contributing to ecological adaptation in in Atlantic herring (Martinez Barrio et al 2016, Han et al 2020). These studies were facilitated by the extremely low genetic differentiation at neutral loci providing an unusually high signal-to-noise ratio in genetic screens. However, the present study has revealed blind spots in these genetic screens, namely the high diversity regions described here, that have likely played a critical role in the evolutionary history of these two species. The high nucleotide diversity within populations at these loci will result in low *F_ST_* estimates also when comparing different populations of Atlantic herring adapted to different environmental conditions, such as those in the marine Atlantic Ocean compared with the brackish Baltic Sea. It is clear that long-read sequencing is required to fully explore the genetic diversity at these loci and how it may contribute to genetic adaptation in herring, and many other vertebrate species.

The Atlantic herring is one of the world’s most abundant vertebrates and a single school may be composed of a billion individuals, making them an attractive target for pathogens. It is very likely that the genetic diversity at immune-related genes described in this study contributes to the genetic defense against pathogens and thus to overall fitness.

## Materials and Methods

### Short-read sequencing data

The Atlantic and Pacific herring samples that were used for population genetic analysis were collected and individually sequenced with Illumina short-read sequencing in our previous studies (Martinez Barrio et al. 2016; Lamichhaney et al. 2017; Han et al. 2020). The samples of Atlantic herring were collected from both the Northeast and Northwest Atlantic Ocean, and the Pacific herring samples were captured close to Vancouver. Information about location and date of sampling, water salinity and spawning season is in Supplementary Table S1, and the procedures of whole genome sequencing and variant detection were described in these references.

### Estimation of population genetic parameters

Population genetic statistics including population differentiation and within and between nucleotide diversity were estimated within non-overlapping 5 kb windows along each chromosome using pixy (Korunes & Samuk 2021). Fixation index (*F_ST_*) was estimated using Weir and Cockerham’s approach (Weir & Cockerham 1984). Average per-site nucleotide differences between all pairs of sequences were calculated within populations (nucleotide diversity, *π*) and between populations (nucleotide divergence, *d_xy_*) within 5 kb windows. Pixy avoids underestimation of nucleotide diversity by using AllSites VCF file which contains both variant and invariant sites. Unbiased estimation of nucleotide diversity is achieved by accounting for missing data in the calculation. Separate files for *F_ST_*, *π* and *d_xy_* and for different chromosomes were combined and windows with a missing value for any of the parameters were excluded from the final file. Windows with the lower 5th percentile of *F_ST_* values between Atlantic and Pacific herring and with the upper 95th percentile of *π* values in each of the two populations were extracted for downstream analyses.

### Genome screening for repetitive elements

We screened the entire Atlantic herring reference genome (Pettersson et al. 2019) for repetitive elements with RepeatMasker (Smit et al. 2013). Then we classified repeat units found by RepeatMasker into superclasses and summarized them over 5 kb windows to match the windows in which diversity parameters were computed. For each window, we calculated the proportion of the window taken up by different superclasses of repeats.

### Gene ontology term overrepresentation analysis

To perform overrepresentation analysis for Gene ontology (GO) terms, we included all protein-coding genes that fully or partially overlapped the windows falling within the lower 5th percentile of *F_ST_* values and the upper 95th percentile of *π* values in each population. Functional annotation for genes residing in these high diversity regions was poor in comparison with the rest of herring genome. To improve functional annotation of herring genes, we employed a Nextflow-based pipeline for functional annotation developed by National Bioinformatics Infrastructure Sweden (NBIS) (https://github.com/NBISweden/pipelines-nextflow). This workflow starts with performing BLAST (Altschul et al. 1990) searches for protein sequences extracted from GFF coordinates against protein database (e-value cut-off was set to 1e-6) and requires UniProt protein fasta file as reference to find the best BLAST matches. Only manually curated proteins (SwissProt proteins) from vertebrates were used. With this approach, it assigns a name to the gene and a description (corresponding to the gene product) to the transcripts. In the next step, it runs InterProScan software package to functionally characterize the genes and assign them functional annotations, including GO terms annotation. NBIS FunctionalAnnotation pipeline in particular helped to identify uncharacterized genes and improved the number of characterized genes from 16,499 to 22,579. In addition, we retrieved GO annotation of orthologues of herring genes from BioMart. We then built the final gene-to-GO map file which links each gene identifier with one or more GO terms. In the final annotation file, 21,345 genes were annotated with a GO term (gene universe), with 587 of them being in the high diversity regions (genes of interest). These genes were used to test overrepresentation of GO terms (17,783, 19,837 and 18,024 genes genome-wide had at least one biological process (BP), molecular function (MF) and cellular component (CC) terms, out of which 270, 503 and 323 genes were in the high diversity regions, respectively).

We used the topGO R package (Alexa & Rahnenfuhrer 2010) for GO term overrepresentation analysis, which provides the possibility to use a custom Gene-to-GO map. The default algorithm, weight01, was used. This algorithm takes the GO topology into account and tests the significance of each GO term depending on its related terms (Alexa et al. 2006). The GO hierarchical structure was read into TopGO from GO.db package. Since the analysis was based on gene count, and no gene score was available, Fisher’s exact test was implemented to assess overrepresentation of GO terms. After removing terms with less than 10 annotated genes, a total of 6,690 terms for BP, 1,319 for MF and 832 for CC were tested and *P* values were adjusted for multiple testing with the Bonferroni method.

### Generation of PacBio long-read data

We generated PacBio HiFi sequencing data from two individuals of Atlantic herring (captured November 25, 2019 in the Celtic Sea). Testis samples were collected and flash-frozen on-site, DNA was extracted using a Circulomics Nanobind Tissue Big DNA Kit (NB-900-701-001), and sequenced using Pacific Biosciences (PacBio) High Fidelity (HiFi) technology. HiFi reads are known to have high accuracy (above 99.8%) and long contiguity (average read length of 13.5 kb) (Wenger et al. 2019) hence we used them to build haploid *de novo* genome assemblies using hifiasm (v0.16.1-r375) (Cheng et al. 2021). The four haplotypes from the PacBio assemblies from the two individuals along with the sequences from the reference assembly were used for further analysis.

We selected three regions from the genome scan to characterize the high diversity regions. As the GO analysis showed a strong enrichment of immune response genes in the high diversity regions, we selected three representative regions that illustrate the complex nature of immune genes including multigene/single gene family and innate/adaptive immune genes. We also considered the contiguity of assembly contigs and gene annotation for selecting these regions. One of the regions encoded for immunoglobulin heavy variable (*IGHV*) multigene family on chromosome 1, and the other two regions were annotated as containing single genes and without an indication of a multigene family on chromosomes 22 and 23, with Ensembl IDs as ENSCHAG00000003891 and ENSCHAG00000015470, respectively. The latter two genes lacked gene name in the Ensembl database, hence to name them, we used nucleotide BLAST (Altschul et al. 1990) to find the most similar gene sequence. CMRF35-like molecule 2 (*CLM2*) and interferon-induced protein with tetratricopeptides 10 (*IFIT10*) were found to be highly similar to ENSCHAG00000003891 and to ENSCHAG00000015470, respectively.

To check for additional copies of these genes, we used nucleotide BLAST (Altschul et al. 1990) and MUMmer (Marçais et al. 2018) aligner using the coding sequences. It resulted in total 40 *IGHV*, four *CLM2*, and two *IFIT10* genes, and a few pseudogenes. Additional *IGHV* genes didn’t have Ensembl IDs, hence these were manually annotated. Additional *CLM2* genes were ENSCHAG00000003799, ENSCHAG00000003851, and ENSCHAG00000003927, while additional *IFIT10* was ENSCHAG00000015478. We named the copies of these additional genes in the order they occurred on the genome – *CLM2A, CLM2B1, CLM2B2*, and *CLM2C* on Chr22, and *IFIT10A* and *IFIT10B* on Chr23.

To annotate these genes on the PacBio assemblies, we used LiftOff (Shumate & Salzberg 2021) with additional “-polish and -copies” parameters. Because *IGHV* is a multigene family, LiftOff algorithm was not successful in annotating all *IGHV* genes hence we did manual curation and used nucleotide BLAST (Altschul et al. 1990) to find homologous sequences using manually annotated reference sequences as queries.

We used gggenomes R package (https://github.com/thackl/gggenomes) to visualize genomic organization of the annotated regions. To calculate accurate *π* for *CLM2* and *IFIT10* genes, we aligned coding sequences using the msa R package (Bodenhofer et al. 2015) and used pegas R package (Paradis 2010) to calculate *π*. The phylogenetic tree was constructed using ape R package (Paradis et al. 2004) and visualized using ggtree (Yu et al. 2017). Homologous genes from the clupeid species *Alosa alosa* (allis shad) were used as an outgroup to root the tree. Genes that were null due to incomplete sequence lengths were excluded for the construction of phylogenetic tree.

## Supplementary Material

Supplementary data are available at *Genome Biology and Evolution* online (http://www.gbe.oxfordjournals.org/).

## Acknowledgments

The project was financially supported by Vetenskapsrådet (2017-02907; to LA) and Knut and Alice Wallenberg Foundation (KAW 2016.0361; to LA). The National Genomics Infrastructure (NGI)/Uppsala Genome Center provided service in massive parallel sequencing and the computational infrastructure was provided by the Swedish National Infrastructure for Computing (SNIC) at UPPMAX partially funded by the Swedish Research Council (2018-05973).

## Data Availability

The raw long-read sequencing data used in this study are deposited in the NCBI Sequence Read Archive (SRA) under the BioProject accession number PRJNA1023520.

