## Supplementary material for "Copy number variation and elevated genetic diversity at immune trait loci in Atlantic and Pacific herring"

### Supplementary material online

#### Supplementary figures

**Figure S1.** Genome-wide distribution of population genetic parameters and repetitive elements in a comparison between Atlantic and Pacific herring across all chromosomes.

**Figure S2.** Correlation matrix for population genetic diversity parameters.

**Figure S3.** Gene ontology subgraphs induced by significant terms in gene set enrichment analysis.

**Figure S4.** Illustrative examples of immune-related gene family clusters at high diversity regions in Atlantic herring reference genome.

#### Supplementary tables

**Table S1.** Samples of Atlantic and Pacific herring used in short-read data analysis.

**Table S2.** Gene ontology term enrichment results for genes overlapping high diversity regions. (Excel file)

**Table S3.** Gene content in high diversity intervals. (Excel file)

**Table S4.** Examples of gene clusters at regions exhibiting low differentiation and high nucleotide diversity in other fishes.

**Table S5.** Pairwise distance matrix for IGHV genes from five haplotypes. (Excel file)

**Table S6.** Nucleotide diversity ($\pi$) based on coding sequences of CLM2 and IFIT10 genes in high diversity regions.

**Figure S1.** Genome-wide distribution of population genetic parameters and repetitive elements in a comparison between Atlantic and Pacific herring for chromosome 1-26. Distribution across chromosome 6 is presented in Figure 1A in the main text. The parameters were estimated within non-overlapping 5 kb windows. Track one to four display population differentiation ($F_{ST}$), intra-population nucleotide diversities in Atlantic ($\pi_{Atlantic}$) and Pacific herring ($\pi_{Pacific}$) and inter-population nucleotide diversity between Atlantic and Pacific herring ($d_{xy}$), respectively. Each dot represents a 5 kb window. Red dots represent windows in the top 5 percentile of $F_{ST}$ and the bottom 5 percentile of $\pi_{Atlantic}$ and $\pi_{Pacific}$. The cumulative proportion of repeats is displayed in the bottom track, and the color code of repeats superfamilies is given below it.

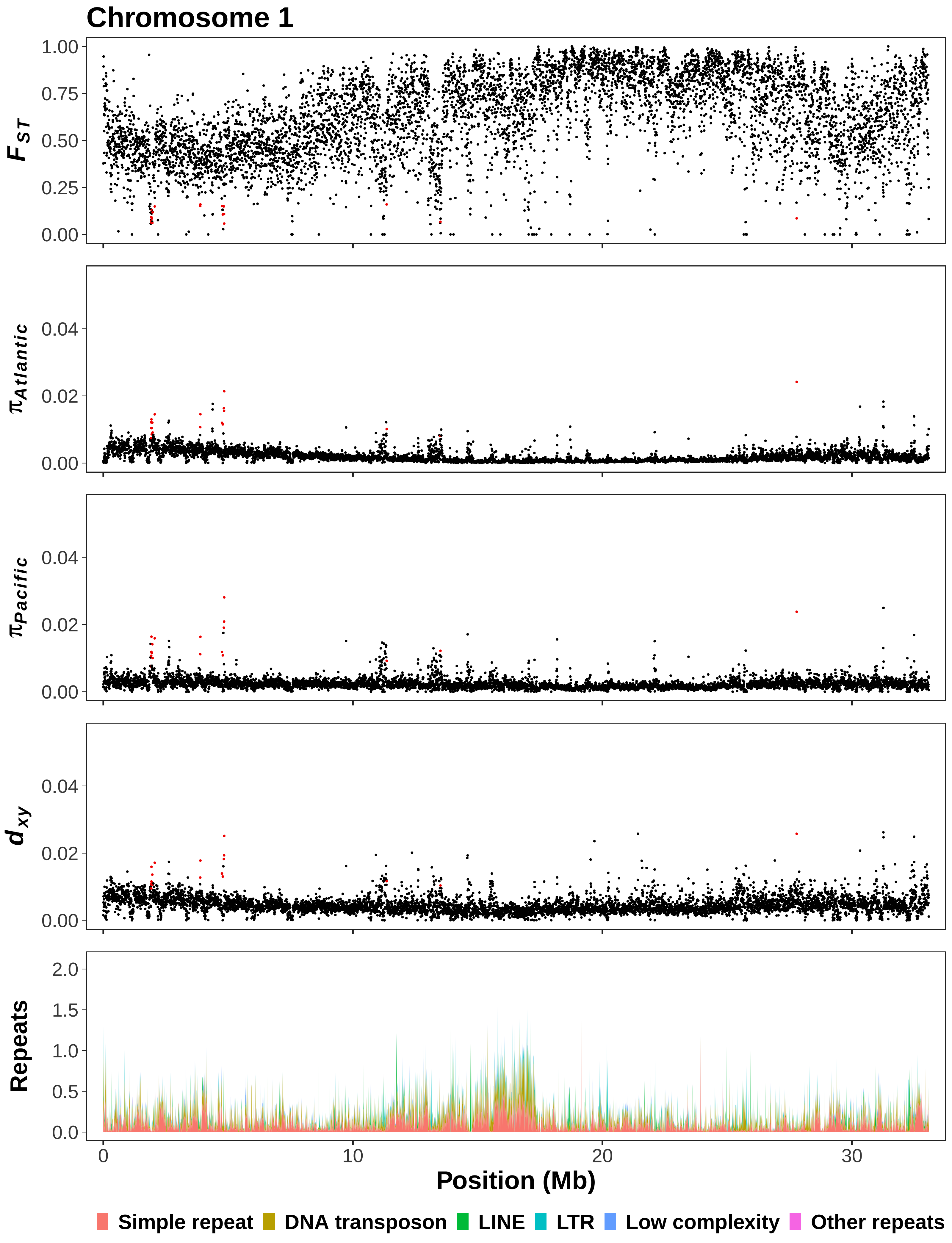

Chromosome 1
F_ST
1.00
0.75
0.50
0.25
0.00
π_Atlantic
0.04
0.02
0.00
π_Pacific
0.04
0.02
0.00
d_xy
0.04
0.02
0.00
Repeats
2.0
1.5
1.0
0.5
0.0
0
10
20
30
Position (Mb)
Simple repeat
DNA transposon
LINE
LTR
Low complexity
Other repeats

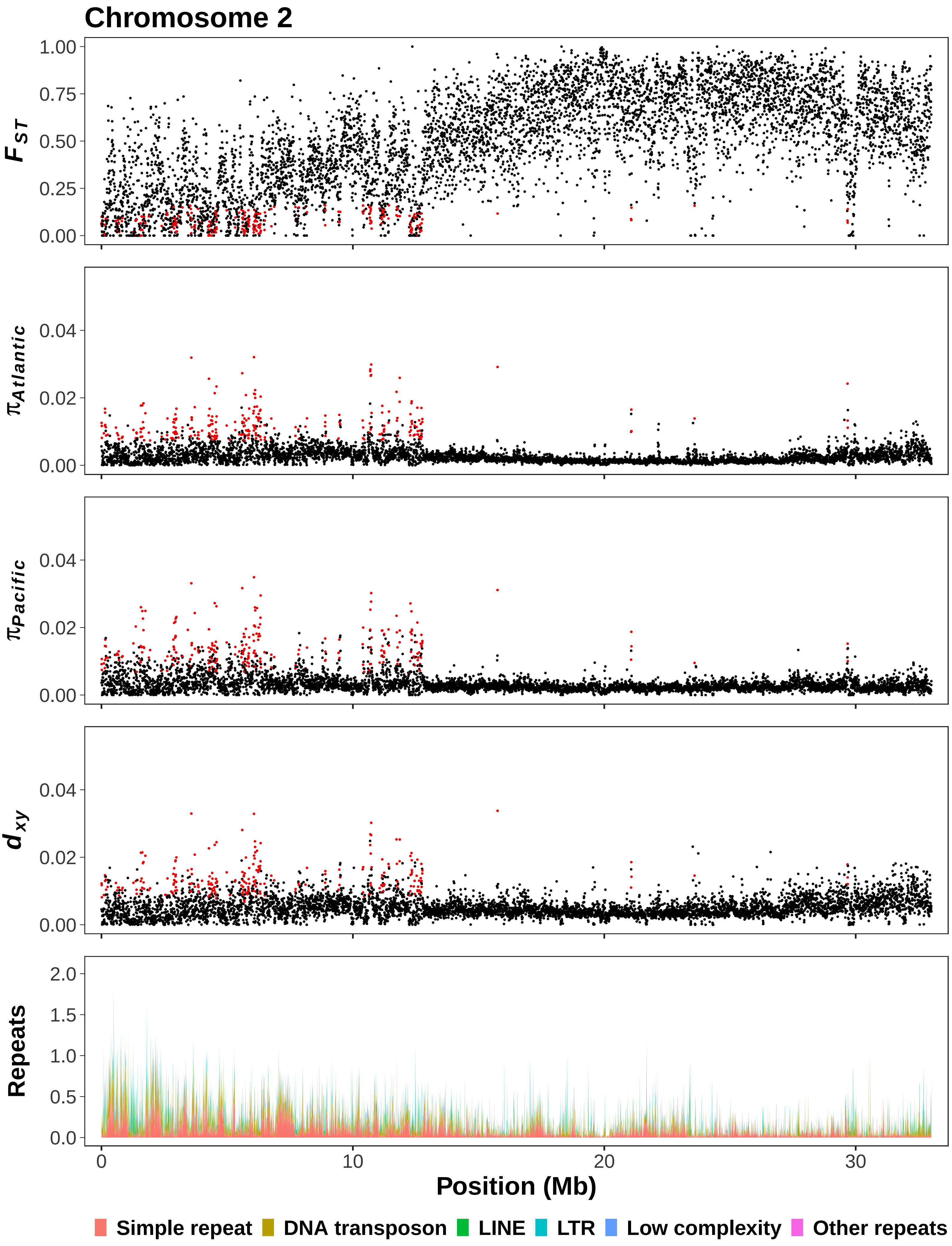

Chromosome 2
$F_{ST}$
1.00
0.75
0.50
0.25
0.00
$\pi_{Atlantic}$
0.04
0.02
0.00
$\pi_{Pacific}$
0.04
0.02
0.00
$d_{xy}$
0.04
0.02
0.00
Repeats
2.0
1.5
1.0
0.5
0.0
0
10
20
30
Position (Mb)
Simple repeat
DNA transposon
LINE
LTR
Low complexity
Other repeats

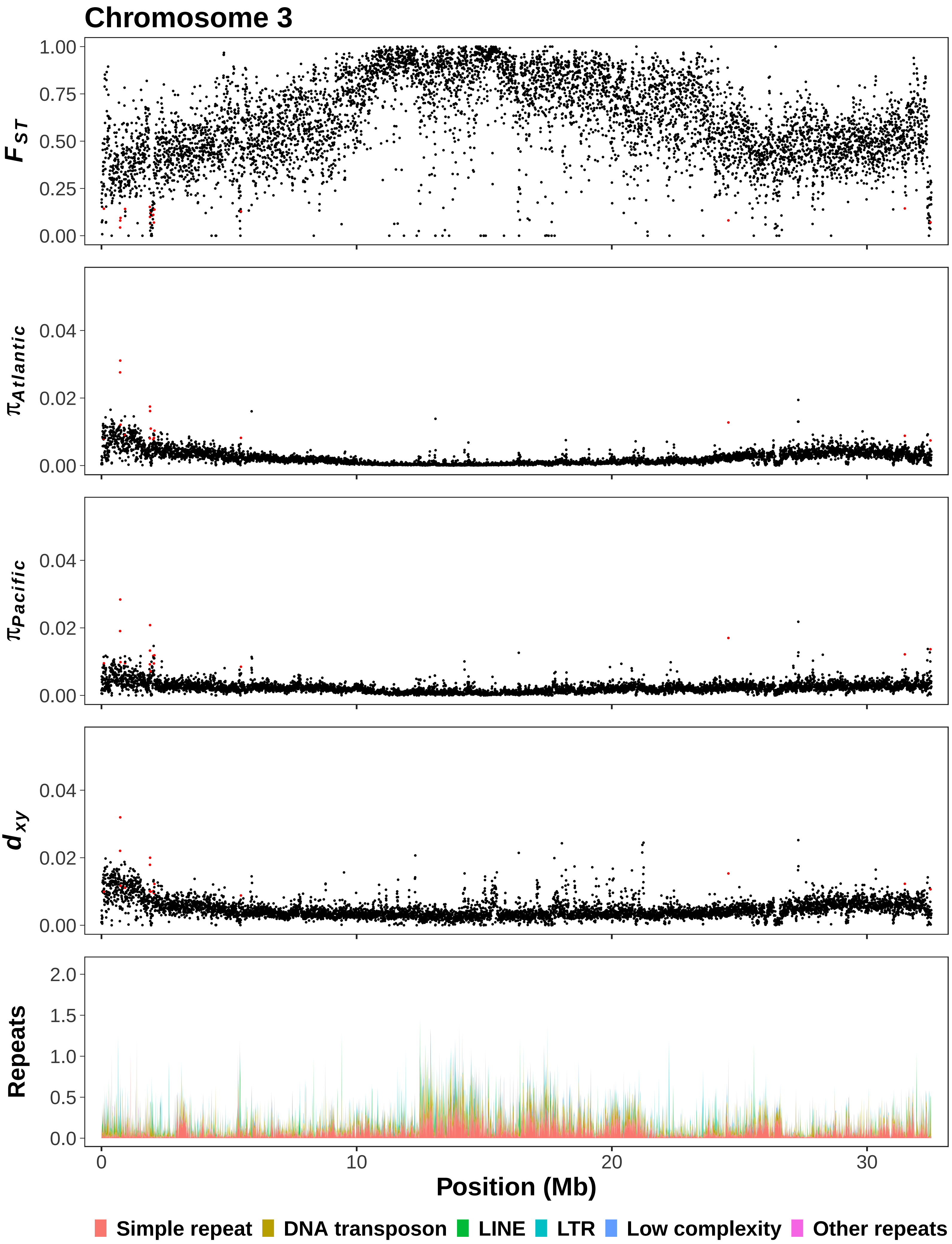

Chromosome 3
$F_{ST}$
1.00
0.75
0.50
0.25
0.00
$\pi_{Atlantic}$
$\pi_{Pacific}$
$d_{xy}$
0.04
0.02
0.00
Repeats
2.0
1.5
1.0
0.5
0.0
0
10
20
30
Position (Mb)
Simple repeat
DNA transposon
LINE
LTR
Low complexity
Other repeats

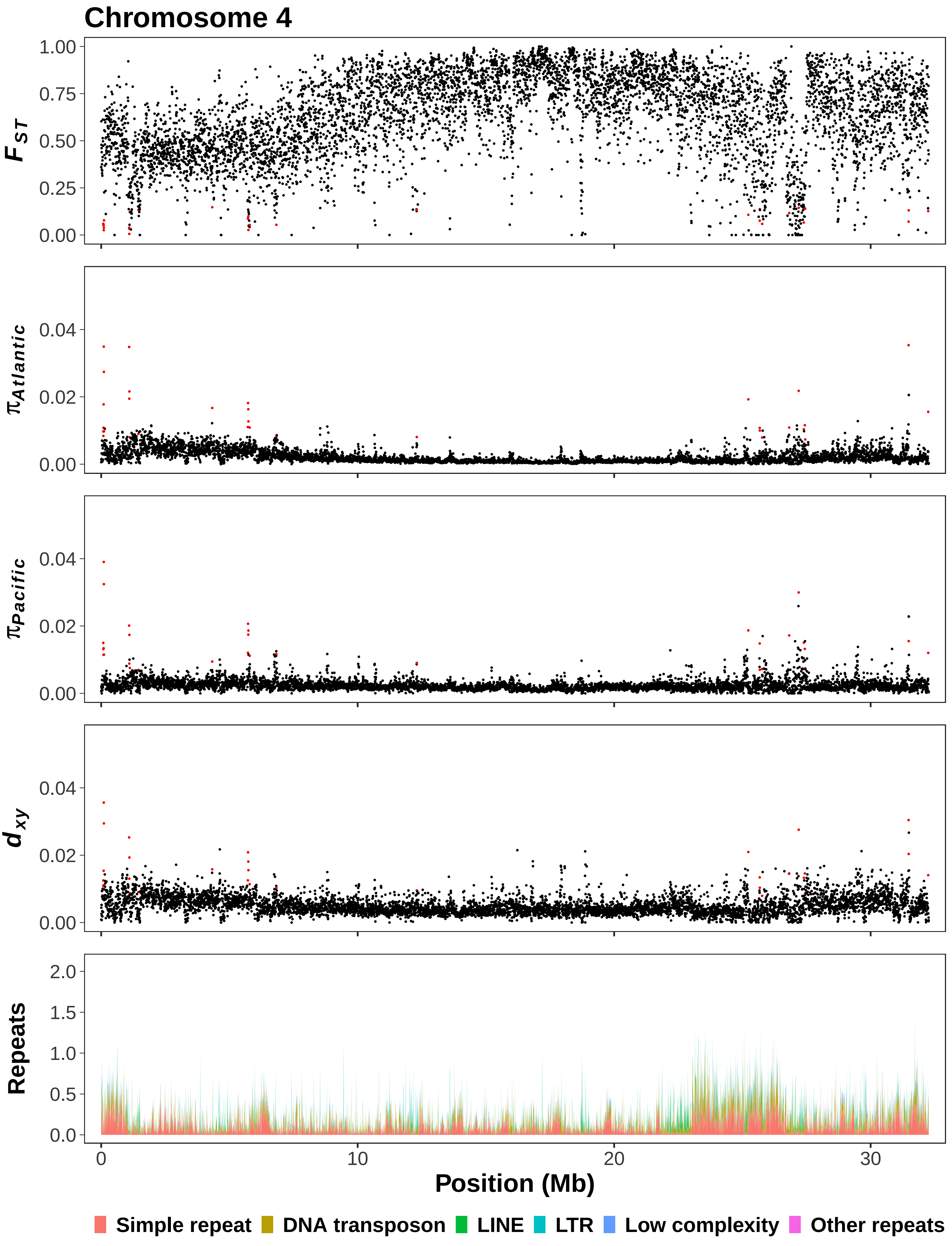

Chromosome 4
$F_{ST}$
1.00
0.75
0.50
0.25
0.00
$\pi_{Atlantic}$
0.04
0.02
0.00
$\pi_{Pacific}$
0.04
0.02
0.00
$d_{xy}$
0.04
0.02
0.00
Repeats
2.0
1.5
1.0
0.5
0.0
0
10
20
30
Position (Mb)
Simple repeat
DNA transposon
LINE
LTR
Low complexity
Other repeats

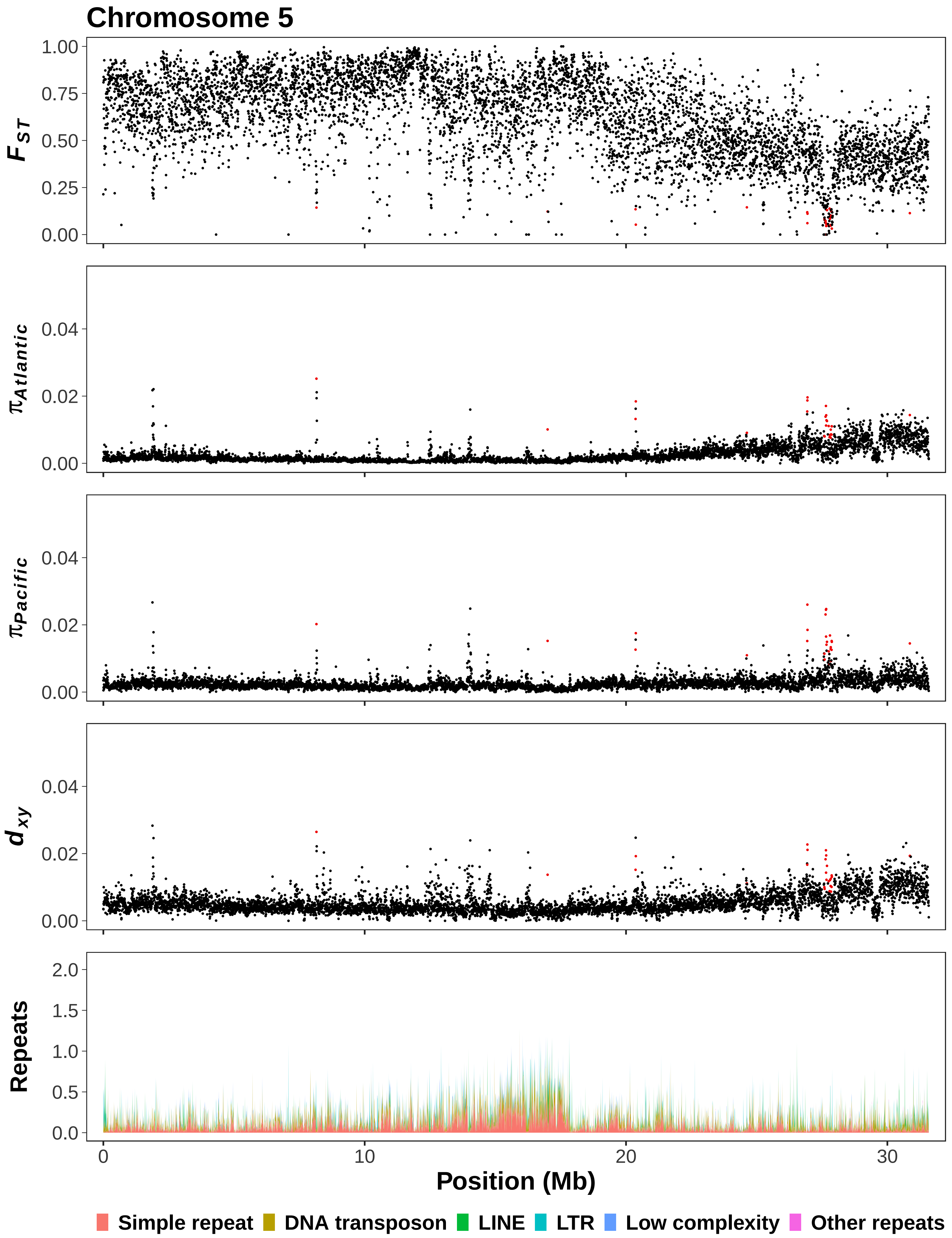

Chromosome 5
$F_{ST}$
1.00
0.75
0.50
0.25
0.00
$\pi_{Atlantic}$
0.04
0.02
0.00
$\pi_{Pacific}$
0.04
0.02
0.00
$d_{xy}$
0.04
0.02
0.00
Repeats
2.0
1.5
1.0
0.5
0.0
0
10
20
30
Position (Mb)
Simple repeat
DNA transposon
LINE
LTR
Low complexity
Other repeats

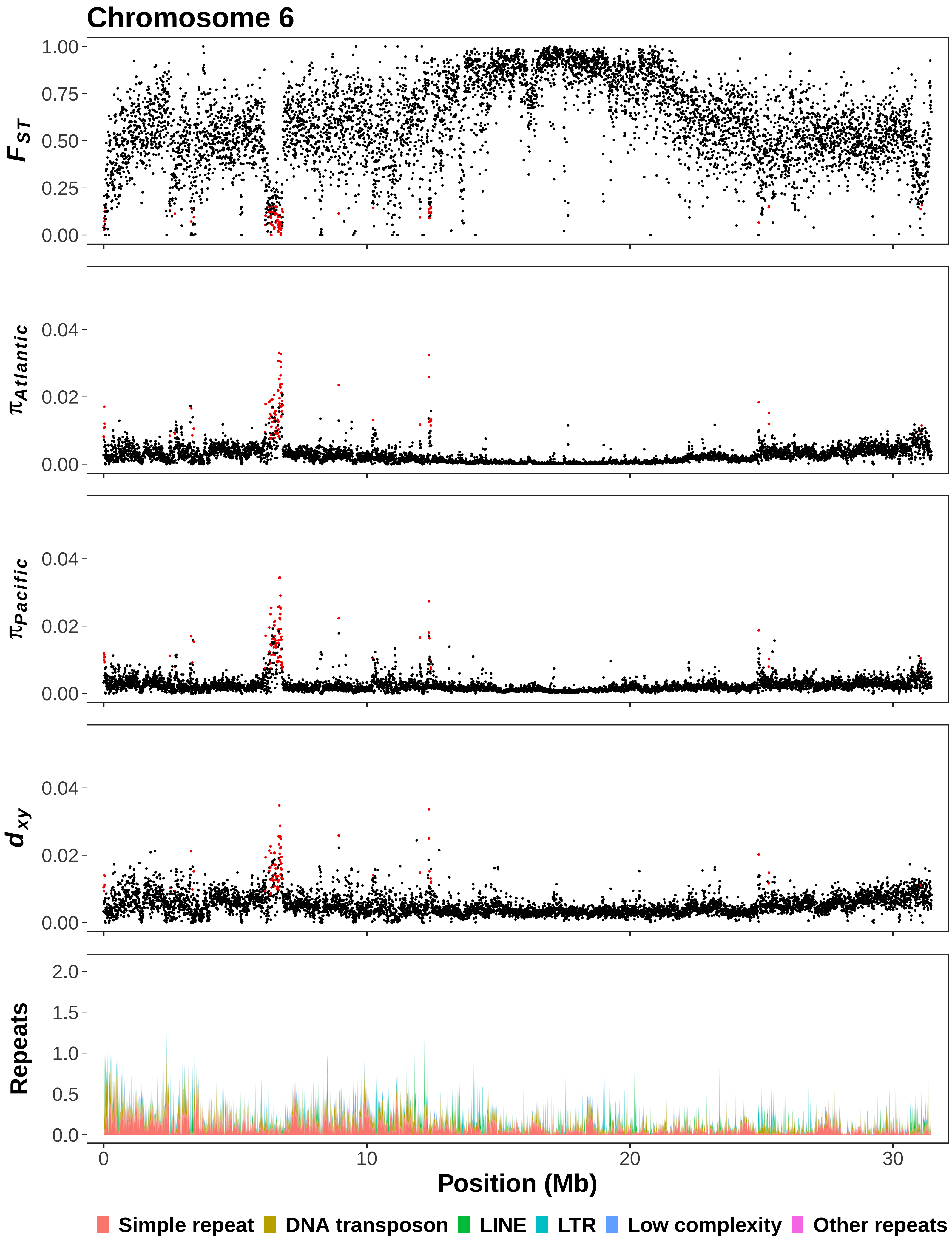

Chromosome 6
$F_{ST}$
1.00
0.75
0.50
0.25
0.00
$\pi_{Atlantic}$
0.04
0.02
0.00
$\pi_{Pacific}$
0.04
0.02
0.00
$d_{xy}$
0.04
0.02
0.00
Repeats
2.0
1.5
1.0
0.5
0.0
0
10
20
30
Position (Mb)
Simple repeat
DNA transposon
LINE
LTR
Low complexity
Other repeats

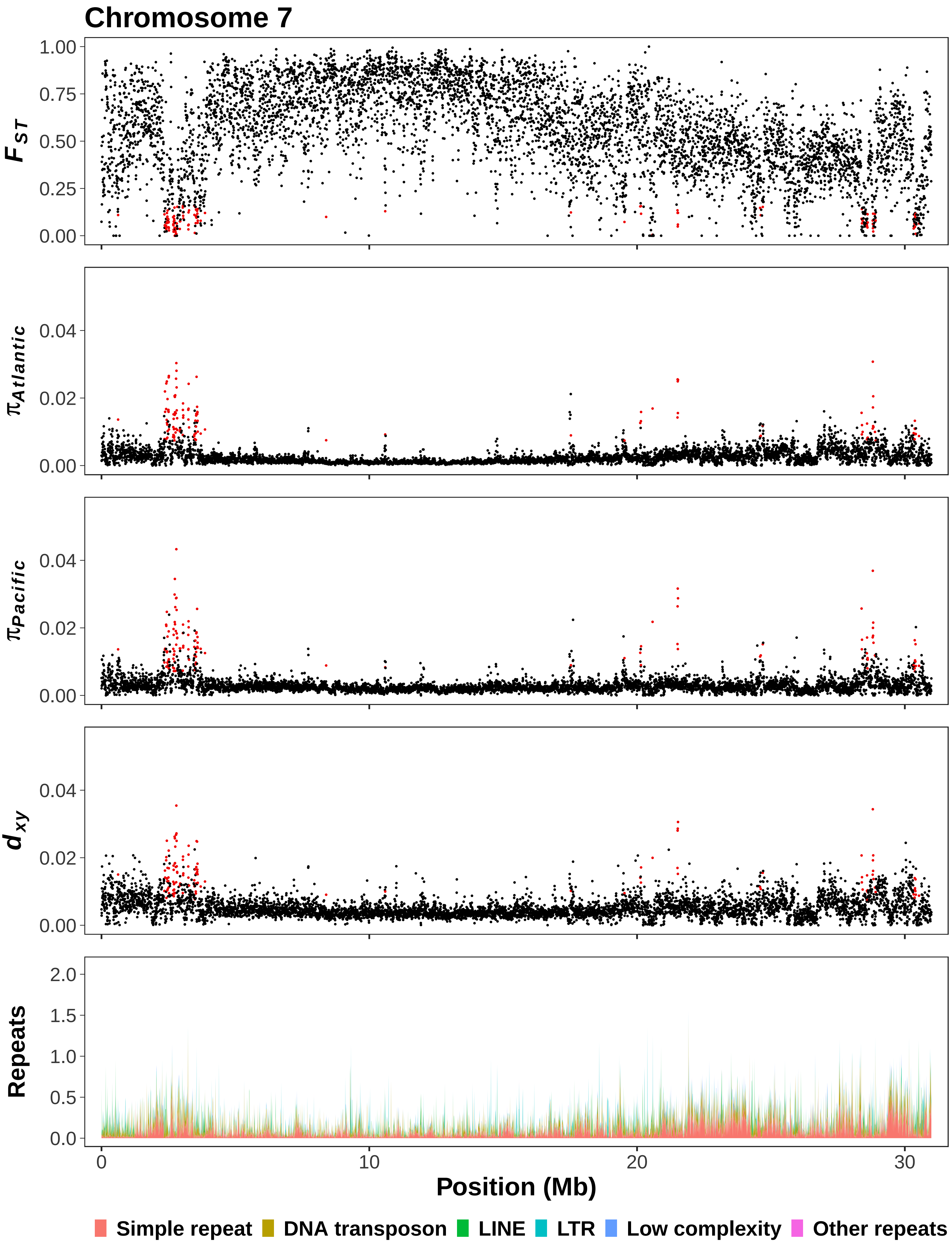

Chromosome 7
$F_{ST}$
1.00
0.75
0.50
0.25
0.00
$\pi_{Atlantic}$
0.04
0.02
0.00
$\pi_{Pacific}$
0.04
0.02
0.00
$d_{xy}$
0.04
0.02
0.00
Repeats
2.0
1.5
1.0
0.5
0.0
0
10
20
30
Position (Mb)
Simple repeat
DNA transposon
LINE
LTR
Low complexity
Other repeats

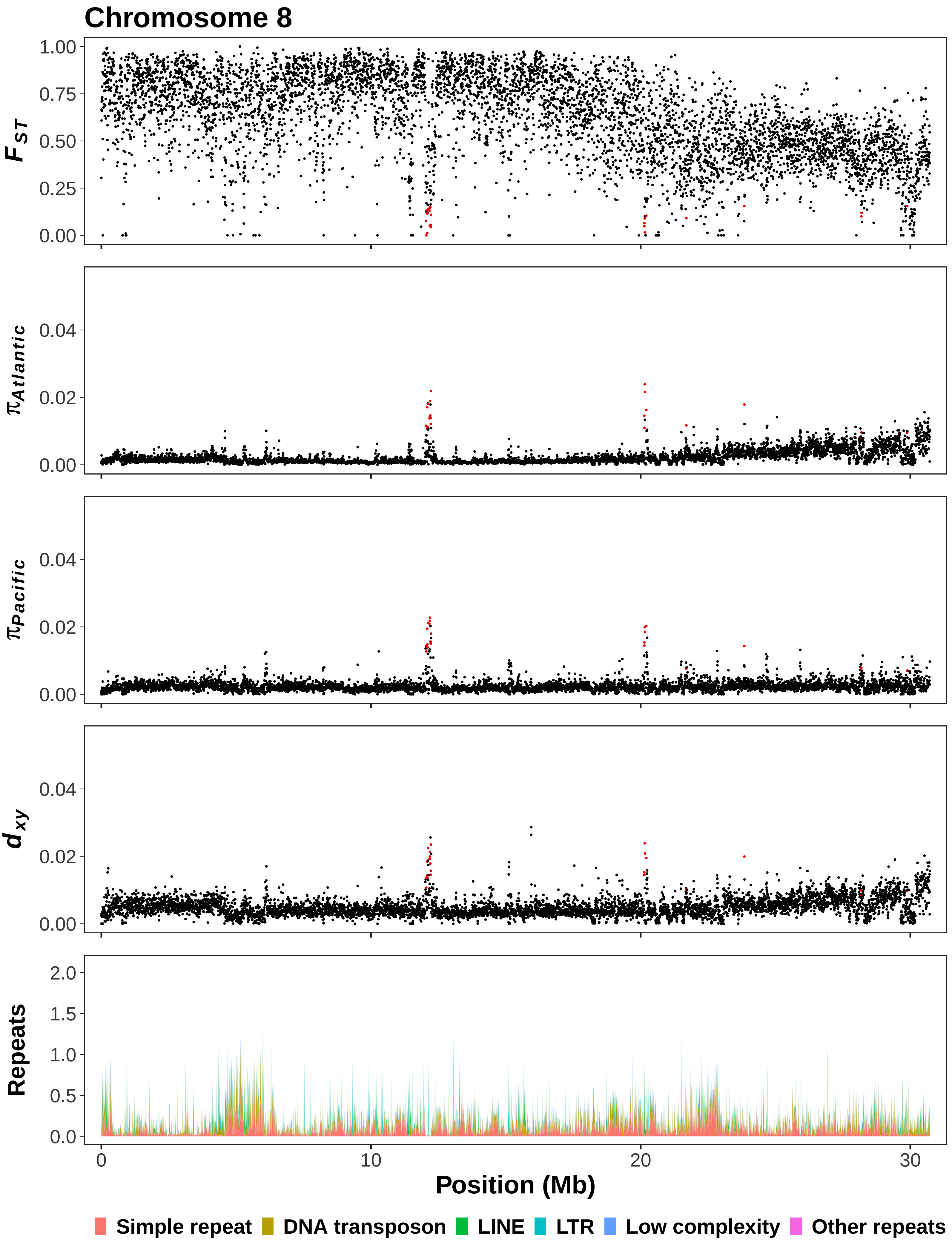

Chromosome 8
$F_{ST}$
1.00
0.75
0.50
0.25
0.00
$\pi_{Atlantic}$
0.04
0.02
0.00
$\pi_{Pacific}$
0.04
0.02
0.00
$d_{xy}$
0.04
0.02
0.00
Repeats
2.0
1.5
1.0
0.5
0.0
0
10
20
30
Position (Mb)
Simple repeat
DNA transposon
LINE
LTR
Low complexity
Other repeats

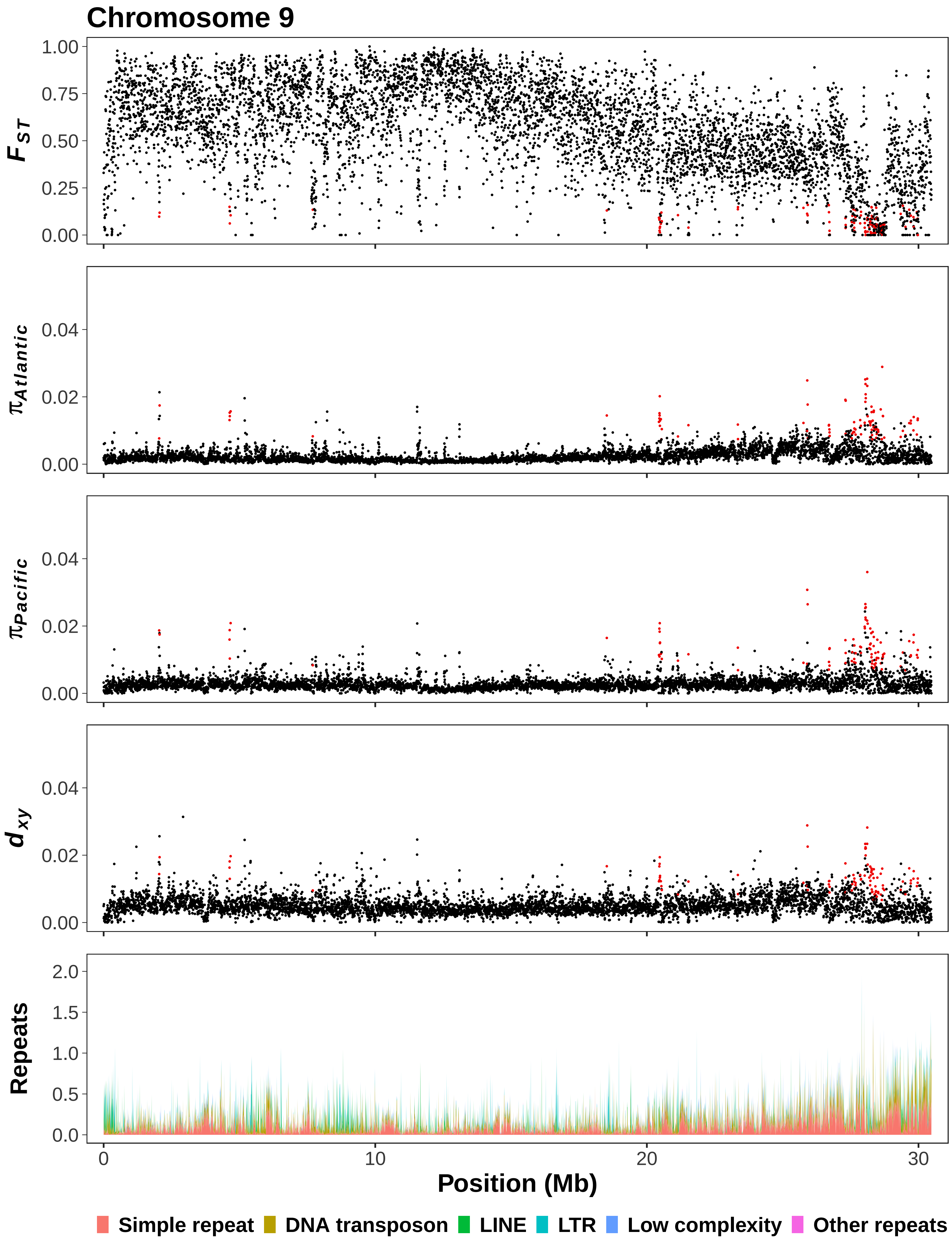

Chromosome 9
$F_{ST}$
1.00
0.75
0.50
0.25
0.00
$\pi_{Atlantic}$
0.04
0.02
0.00
$\pi_{Pacific}$
0.04
0.02
0.00
$d_{xy}$
0.04
0.02
0.00
Repeats
2.0
1.5
1.0
0.5
0.0
0
10
20
30
Position (Mb)
Simple repeat
DNA transposon
LINE
LTR
Low complexity
Other repeats

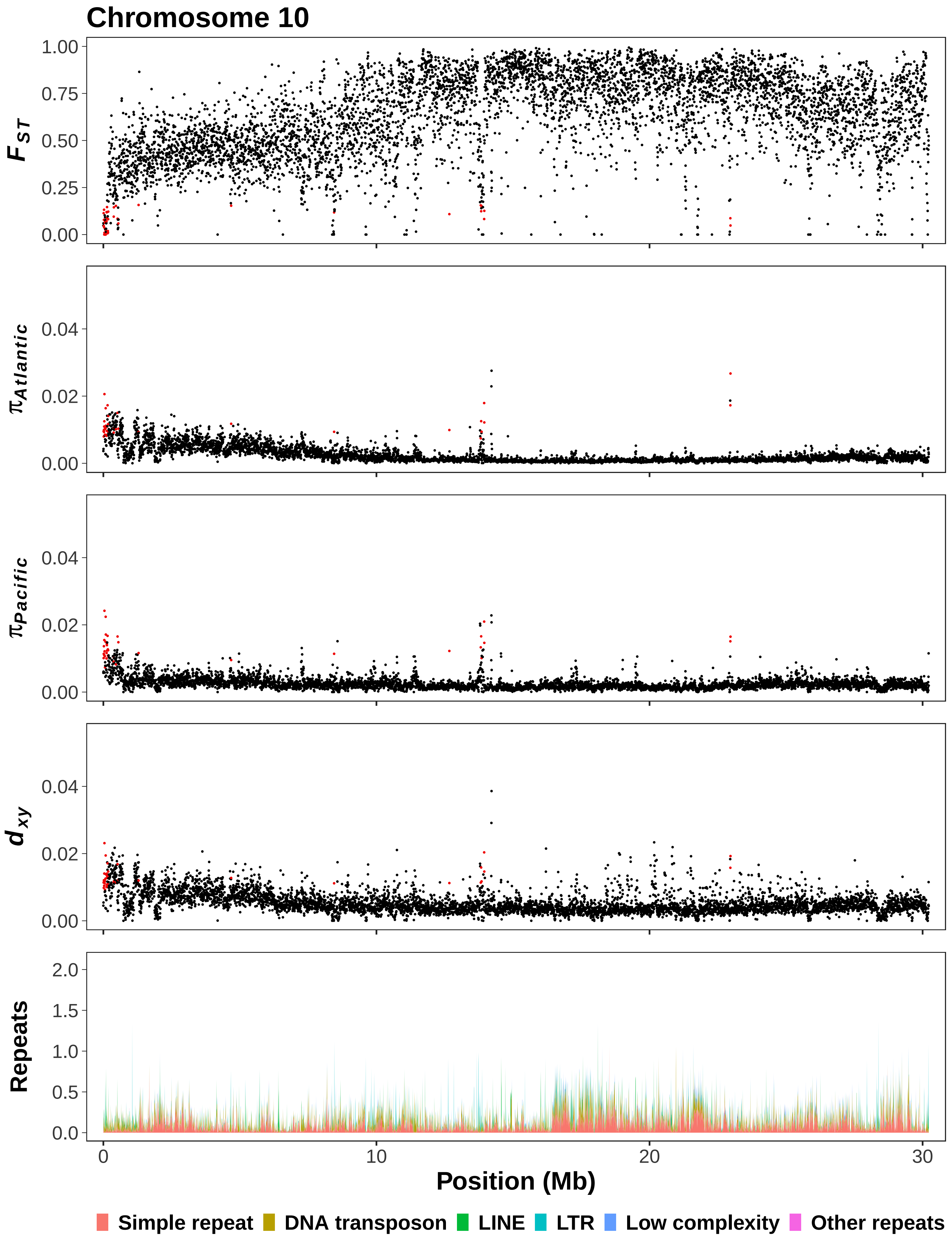

Chromosome 10
$F_{ST}$
1.00
0.75
0.50
0.25
0.00
$\pi_{Atlantic}$
0.04
0.02
0.00
$\pi_{Pacific}$
$d_{xy}$
Repeats
2.0
1.5
1.0
0.5
0.0
0
10
20
30
Position (Mb)
Simple repeat
DNA transposon
LINE
LTR
Low complexity
Other repeats

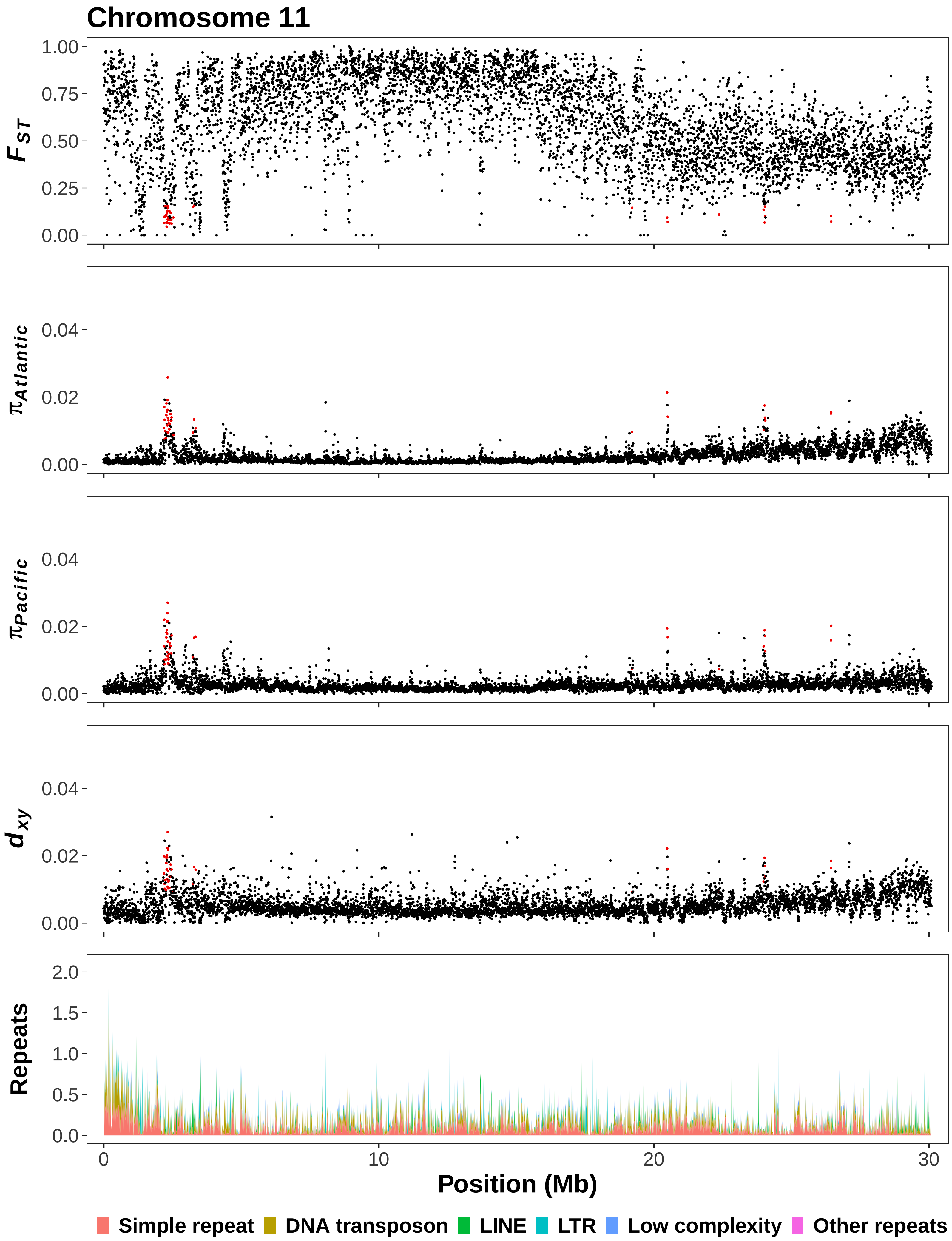

Chromosome 11
$F_{ST}$
1.00
0.75
0.50
0.25
0.00
$\pi_{Atlantic}$
0.04
0.02
0.00
$\pi_{Pacific}$
0.04
0.02
0.00
$d_{xy}$
0.04
0.02
0.00
Repeats
2.0
1.5
1.0
0.5
0.0
0
10
20
30
Position (Mb)
Simple repeat
DNA transposon
LINE
LTR
Low complexity
Other repeats

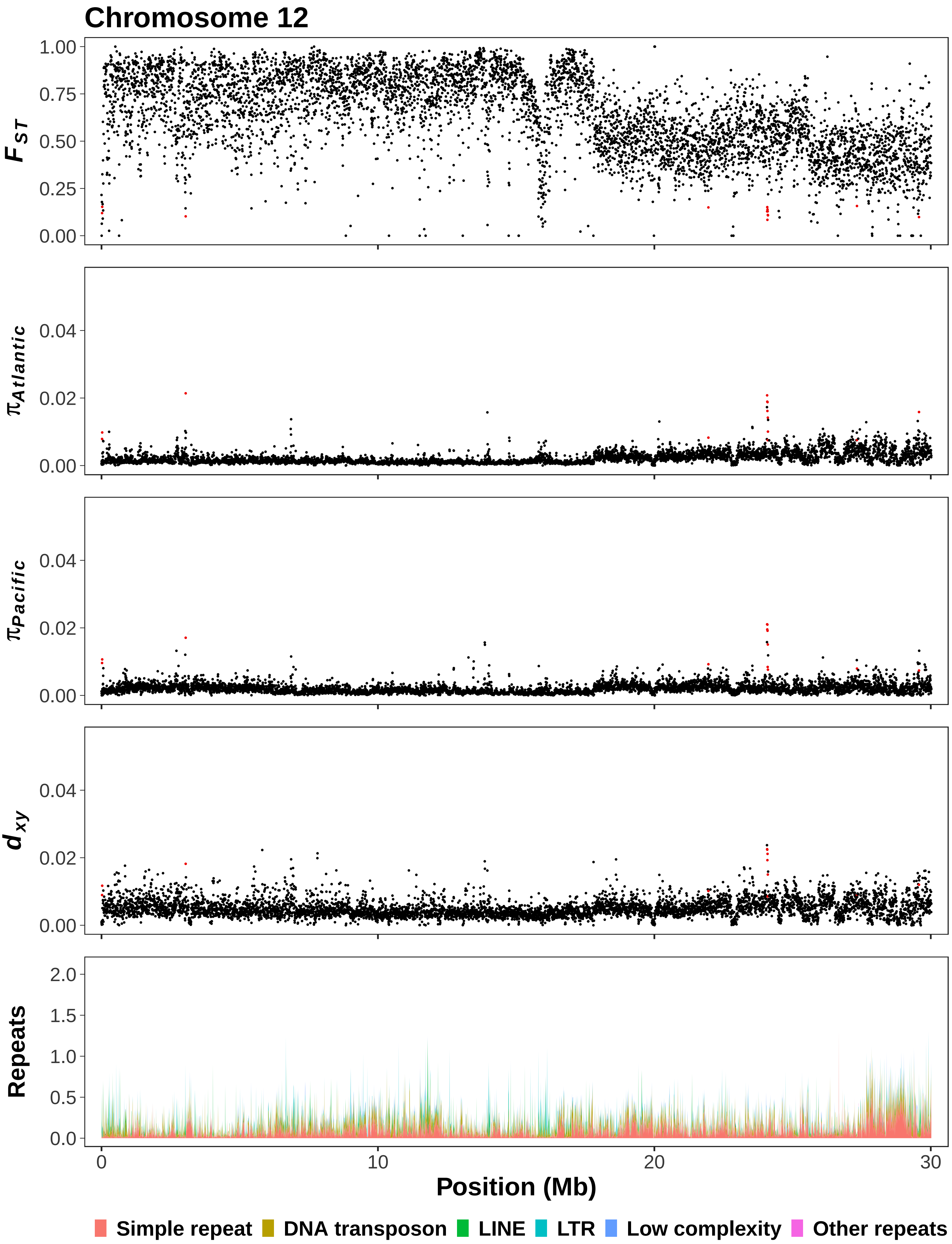

Chromosome 12
$F_{ST}$
1.00
0.75
0.50
0.25
0.00
$\pi_{Atlantic}$
0.04
0.02
0.00
$\pi_{Pacific}$
0.04
0.02
0.00
$d_{xy}$
0.04
0.02
0.00
Repeats
2.0
1.5
1.0
0.5
0.0
0
10
20
30
Position (Mb)
Simple repeat
DNA transposon
LINE
LTR
Low complexity
Other repeats

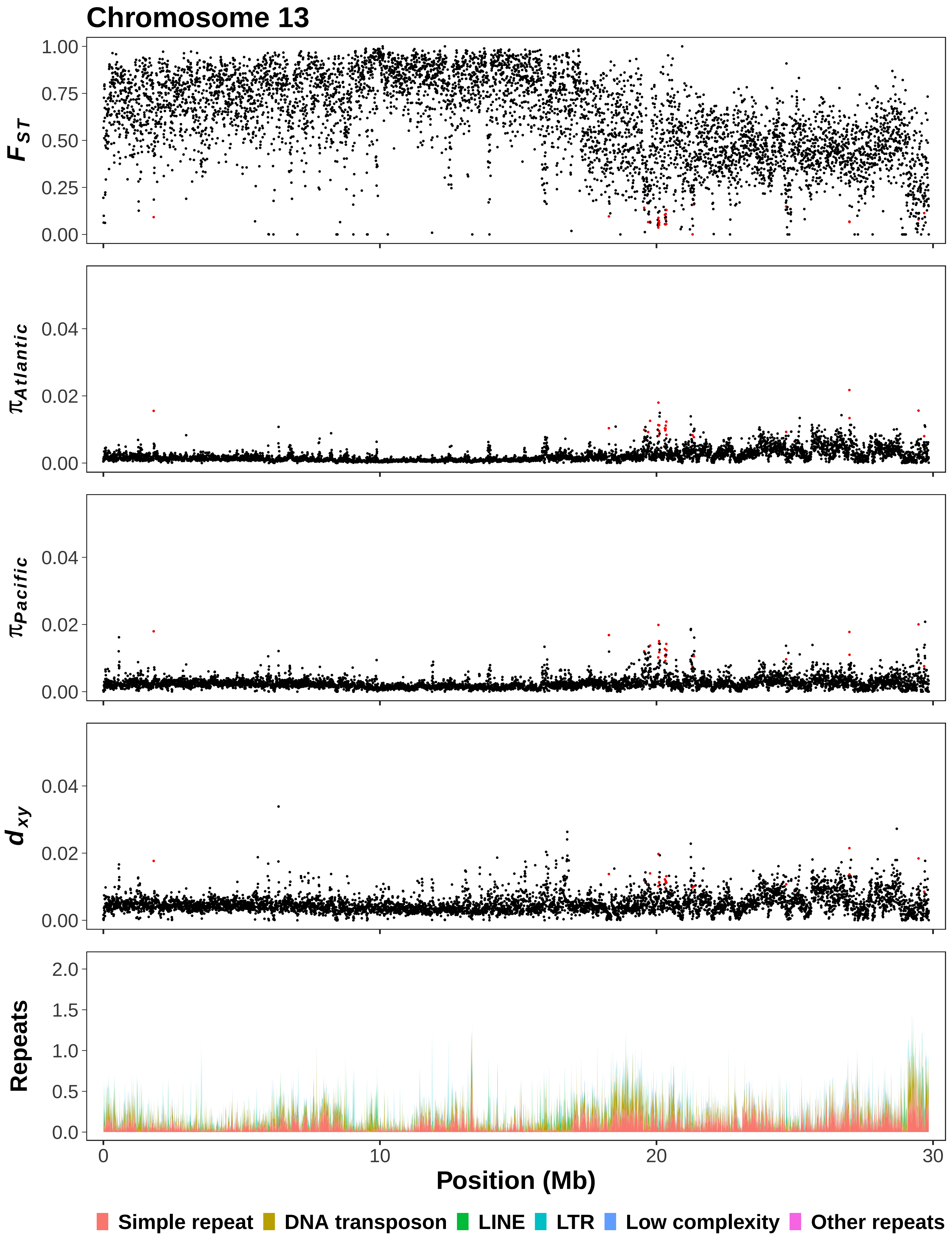

Chromosome 13
$F_{ST}$
1.00
0.75
0.50
0.25
0.00
$\pi_{Atlantic}$
0.04
0.02
0.00
$\pi_{Pacific}$
0.04
0.02
0.00
$d_{xy}$
0.04
0.02
0.00
Repeats
2.0
1.5
1.0
0.5
0.0
0
10
20
30
Position (Mb)
Simple repeat
DNA transposon
LINE
LTR
Low complexity
Other repeats

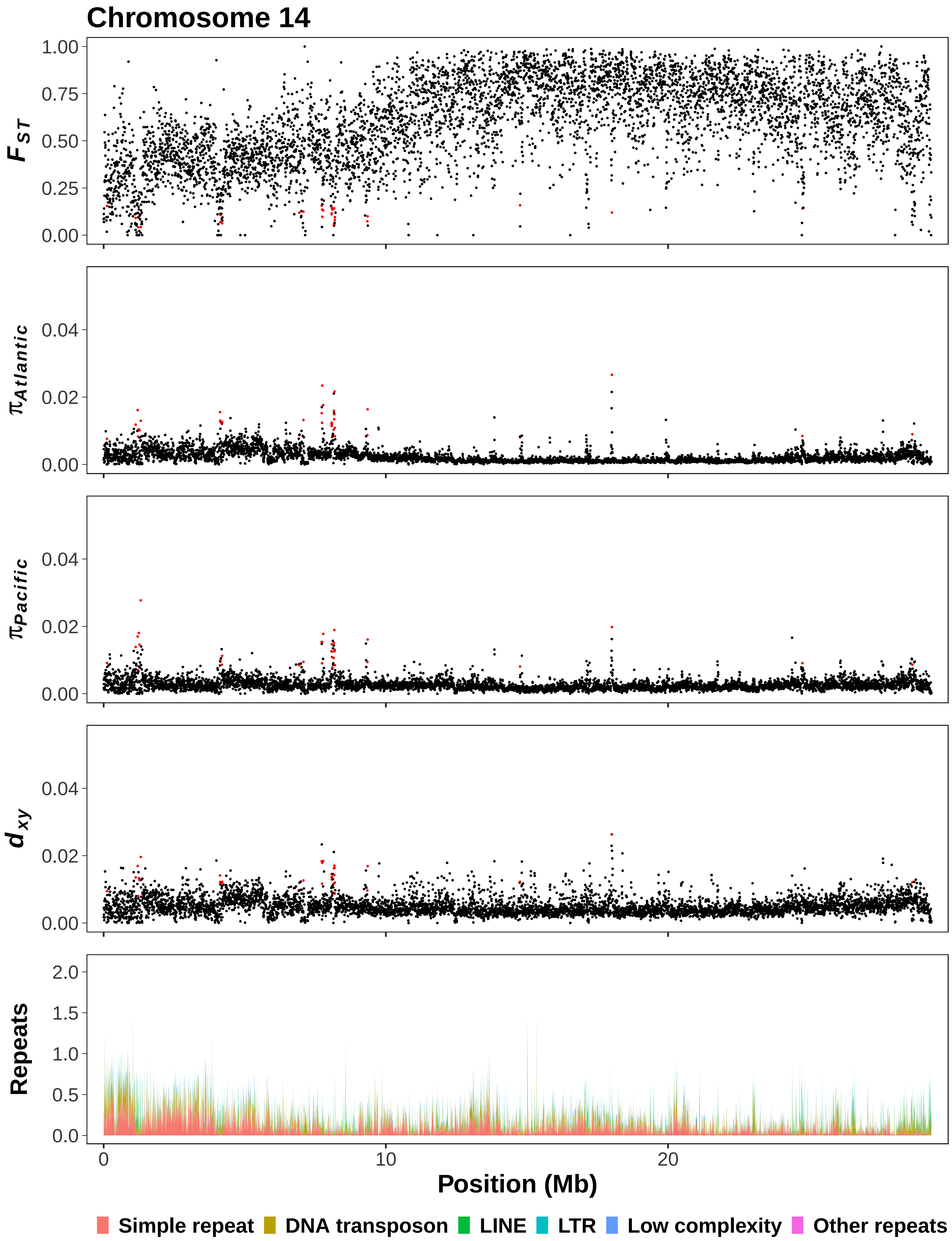

Chromosome 14
$F_{ST}$
$\pi_{Atlantic}$
$\pi_{Pacific}$
$d_{xy}$
Repeats
Position (Mb)
Simple repeat
DNA transposon
LINE
LTR
Low complexity
Other repeats

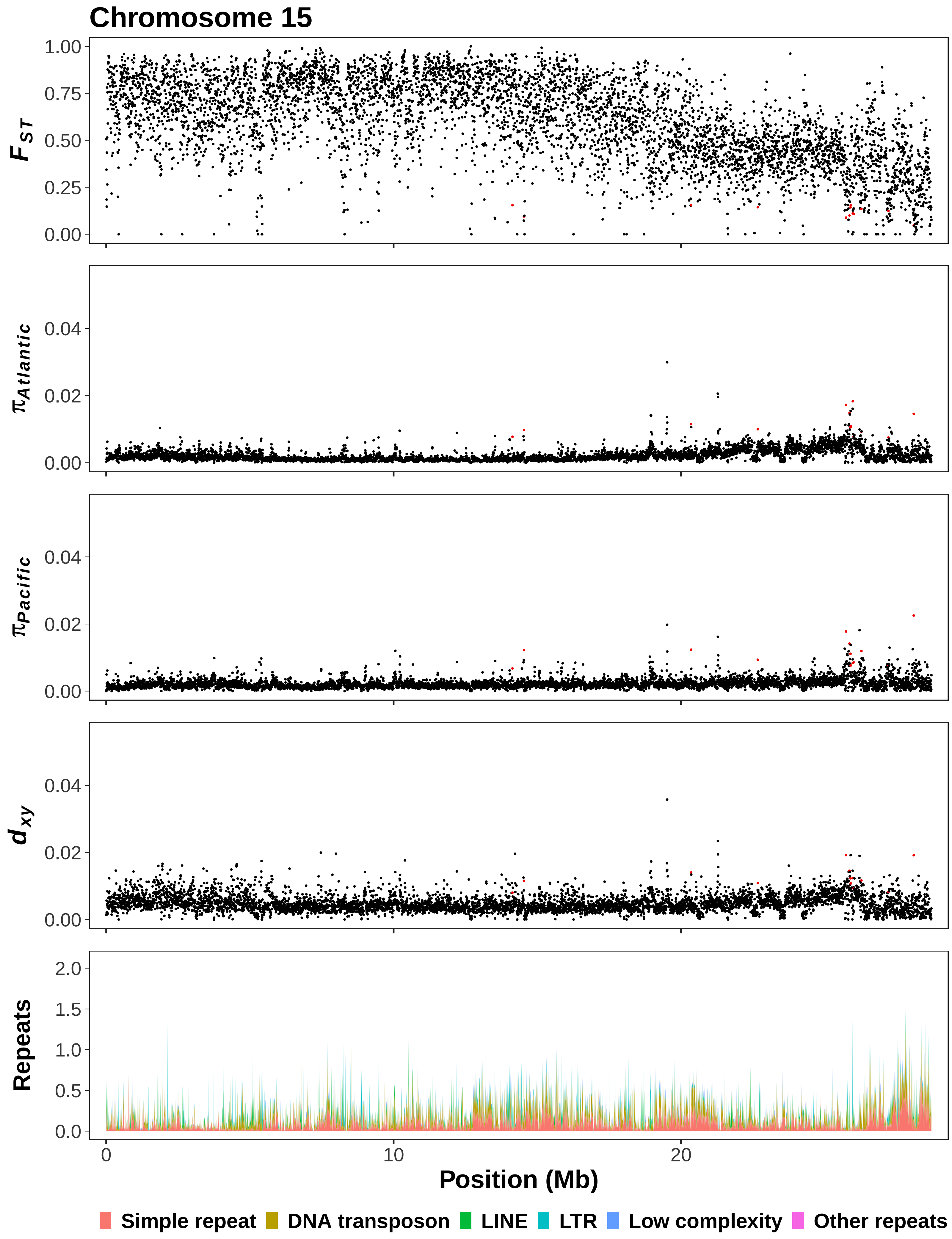

Chromosome 15
$F_{ST}$
1.00
0.75
0.50
0.25
0.00
$\pi_{Atlantic}$
0.04
0.02
0.00
$\pi_{Pacific}$
0.04
0.02
0.00
$d_{xy}$
0.04
0.02
0.00
Repeats
2.0
1.5
1.0
0.5
0.0
0
10
20
Position (Mb)
Simple repeat
DNA transposon
LINE
LTR
Low complexity
Other repeats

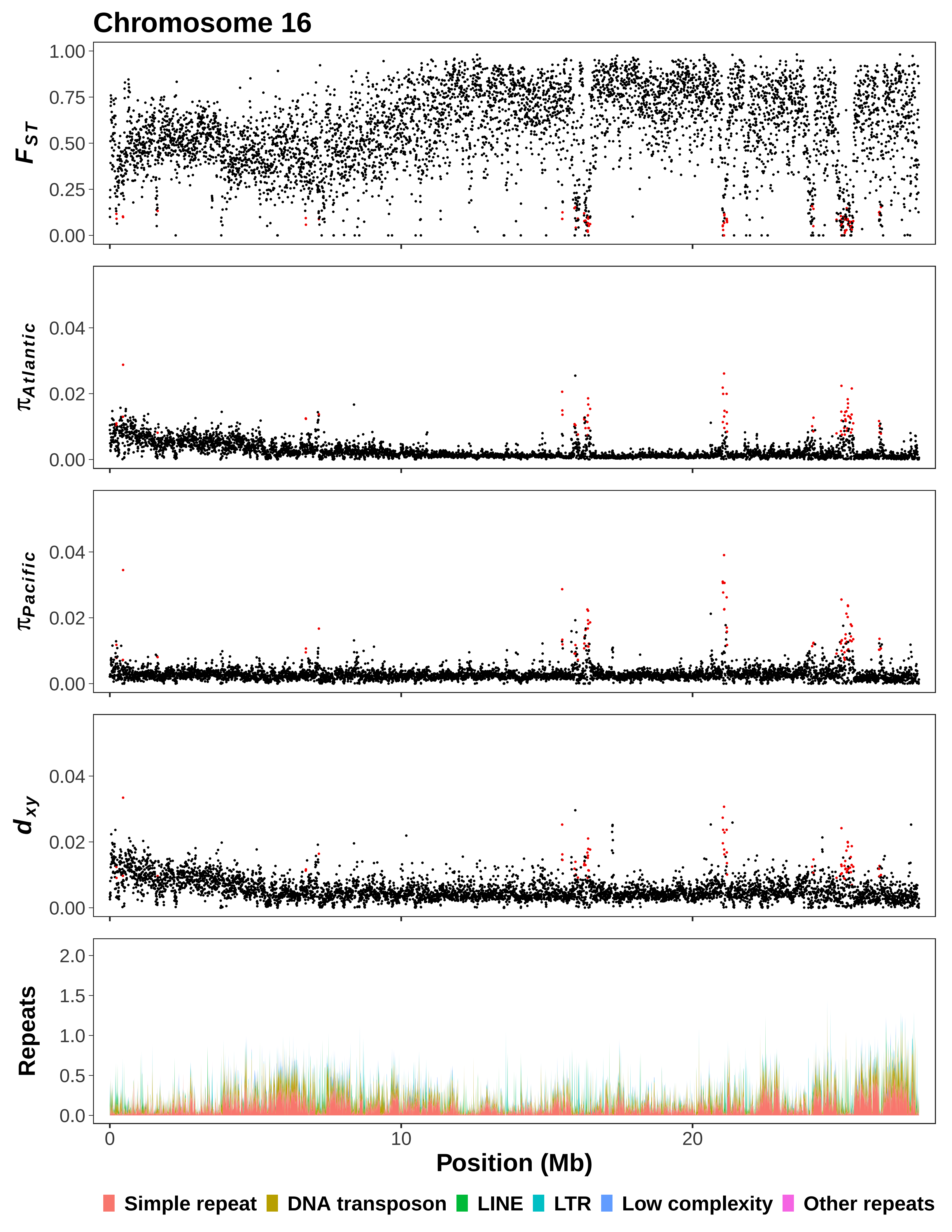
Chromosome 16
F_ST
π_Atlantic
π_Pacific
d_xy
Repeats
Position (Mb)
Simple repeat
DNA transposon
LINE
LTR
Low complexity
Other repeats

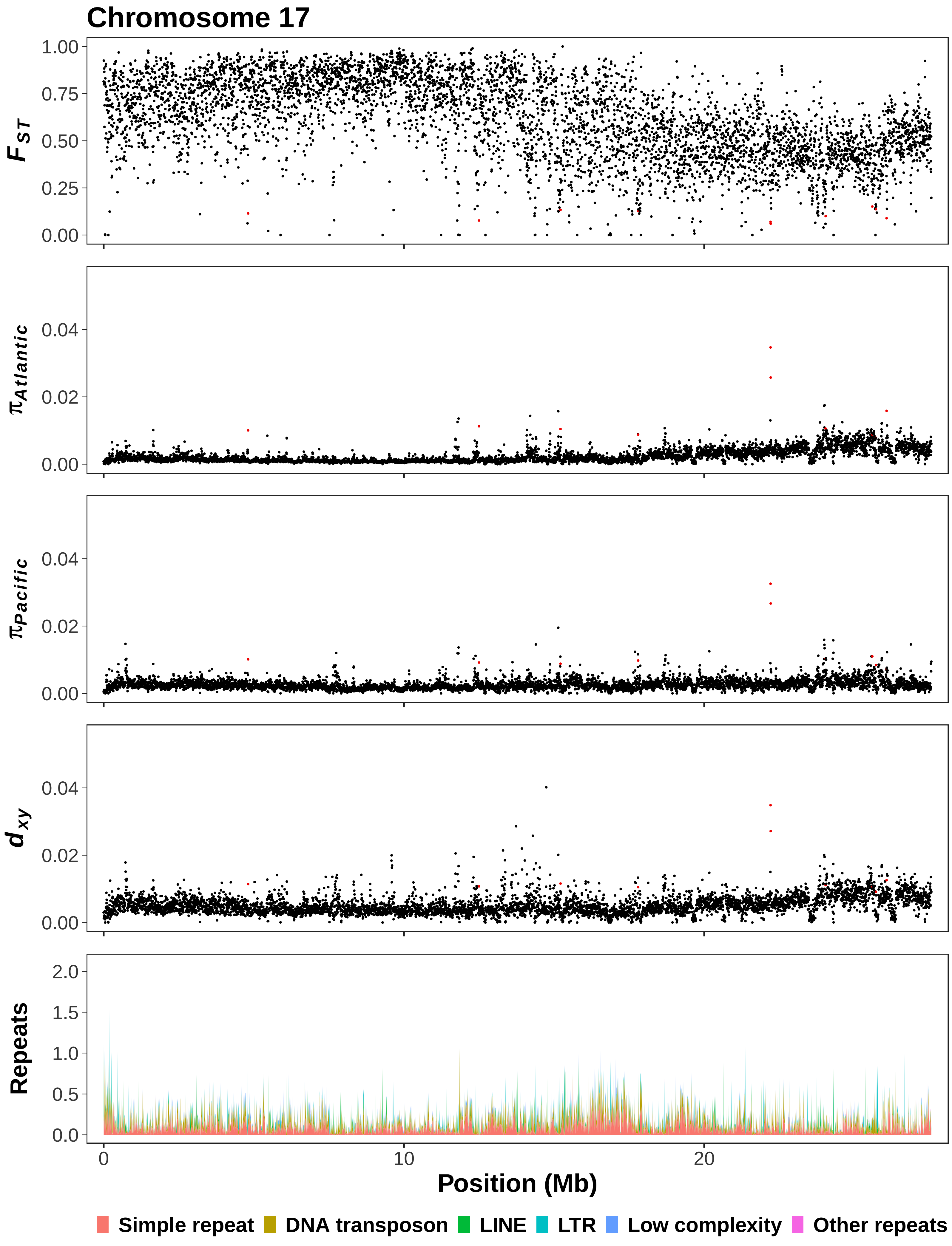

Chromosome 17
F_ST
1.00
0.75
0.50
0.25
0.00
π_Atlantic
0.04
0.02
0.00
π_Pacific
0.04
0.02
0.00
d_xy
0.04
0.02
0.00
Repeats
2.0
1.5
1.0
0.5
0.0
0
10
20
Position (Mb)
Simple repeat
DNA transposon
LINE
LTR
Low complexity
Other repeats

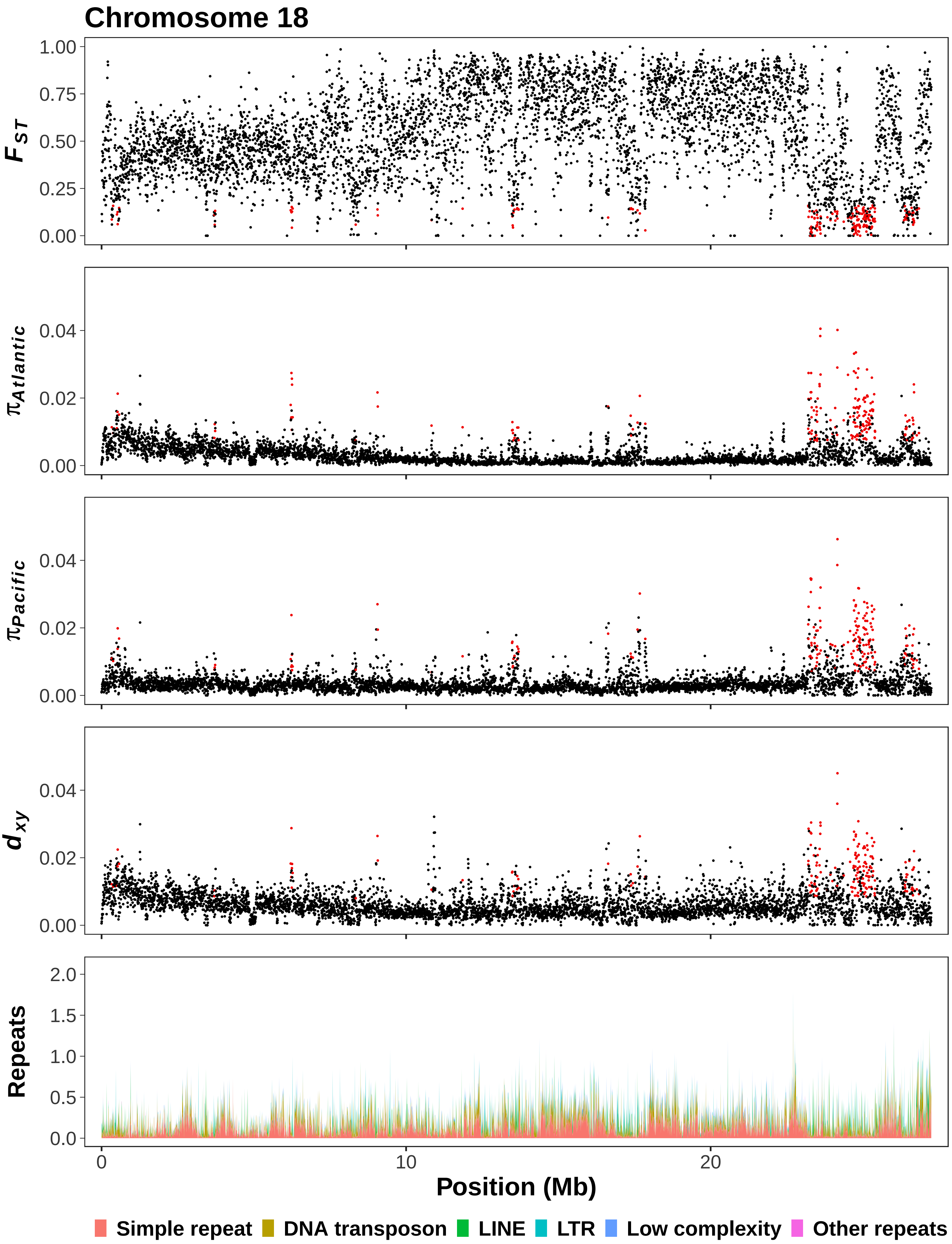

Chromosome 18
F_ST
π_Atlantic
π_Pacific
d_xy
Repeats
1.00
0.75
0.50
0.25
0.00
0.04
0.02
0.00
2.0
1.5
1.0
0.5
0.0
0
10
20
Position (Mb)
Simple repeat
DNA transposon
LINE
LTR
Low complexity
Other repeats

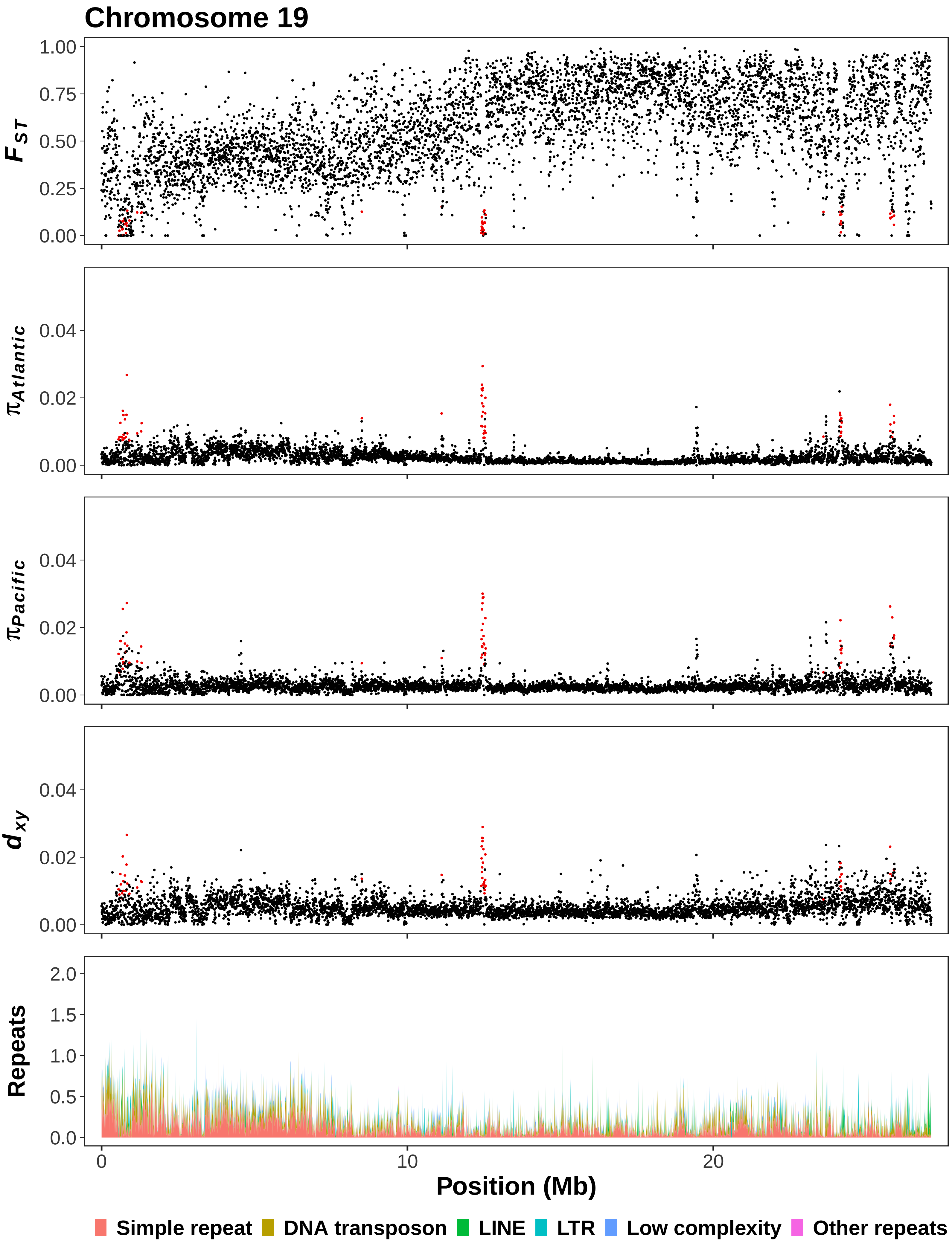

Chromosome 19
$F_{ST}$
1.00
0.75
0.50
0.25
0.00
$\pi_{Atlantic}$
0.04
0.02
0.00
$\pi_{Pacific}$
0.04
0.02
0.00
$d_{xy}$
0.04
0.02
0.00
Repeats
2.0
1.5
1.0
0.5
0.0
0
10
20
Position (Mb)
Simple repeat
DNA transposon
LINE
LTR
Low complexity
Other repeats

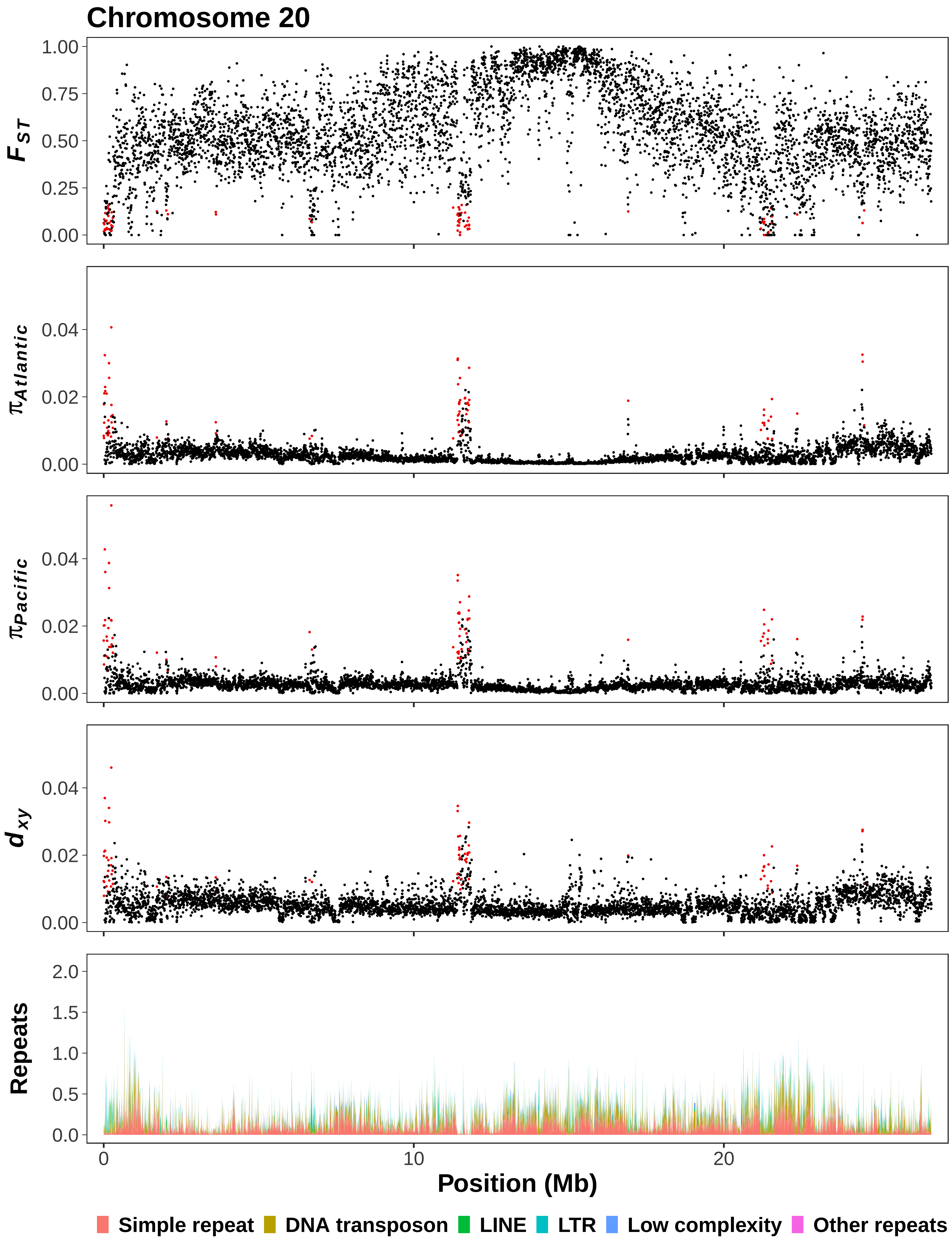

Chromosome 20
$F_{ST}$
1.00
0.75
0.50
0.25
0.00
$\pi_{Atlantic}$
0.04
0.02
0.00
$\pi_{Pacific}$
0.04
0.02
0.00
$d_{xy}$
0.04
0.02
0.00
Repeats
2.0
1.5
1.0
0.5
0.0
0
10
20
Position (Mb)
Simple repeat
DNA transposon
LINE
LTR
Low complexity
Other repeats

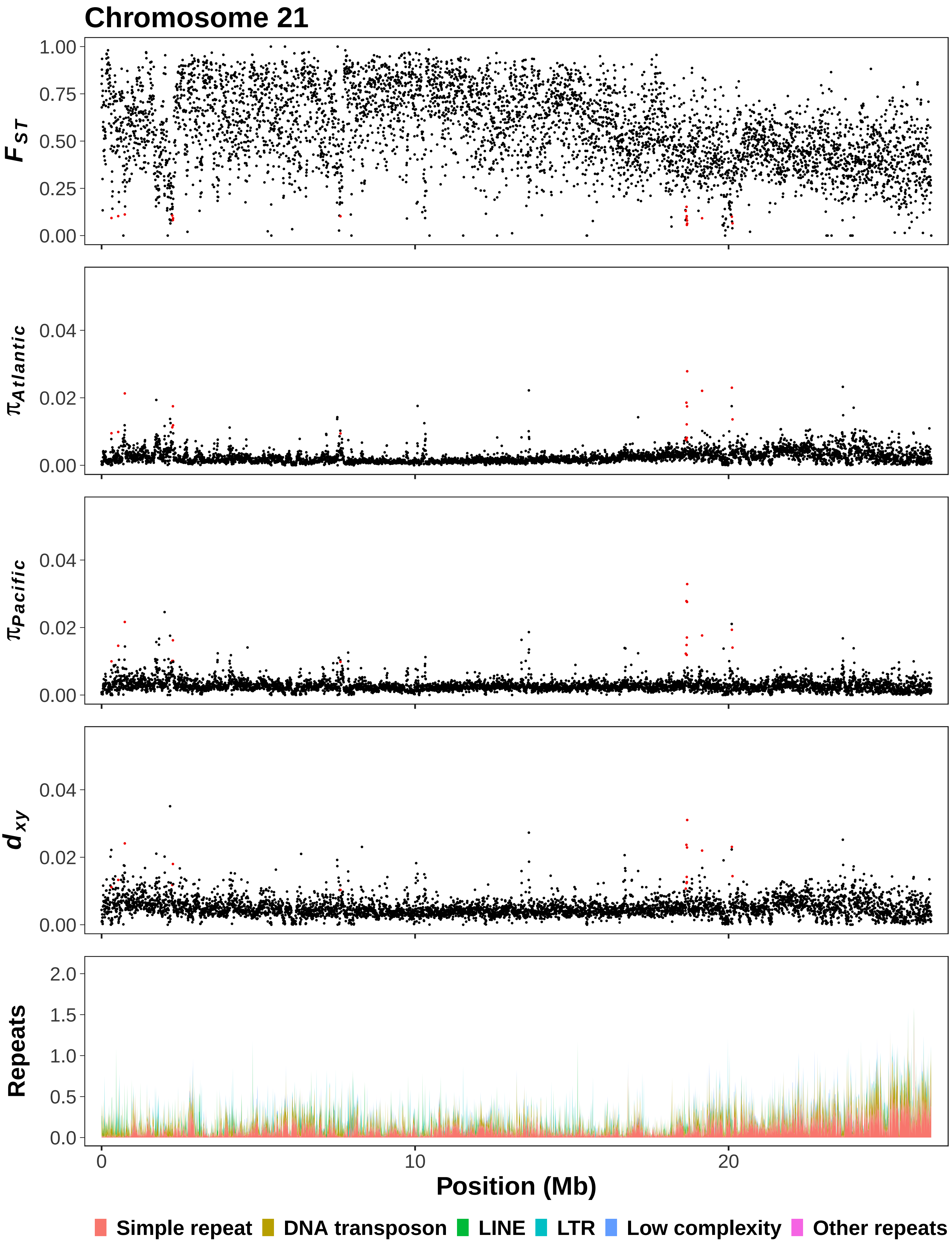

Chromosome 21
$F_{ST}$
1.00
0.75
0.50
0.25
0.00
$\pi_{Atlantic}$
$\pi_{Pacific}$
$d_{xy}$
0.04
0.02
0.00
Repeats
2.0
1.5
1.0
0.5
0.0
0
10
20
Position (Mb)
Simple repeat
DNA transposon
LINE
LTR
Low complexity
Other repeats

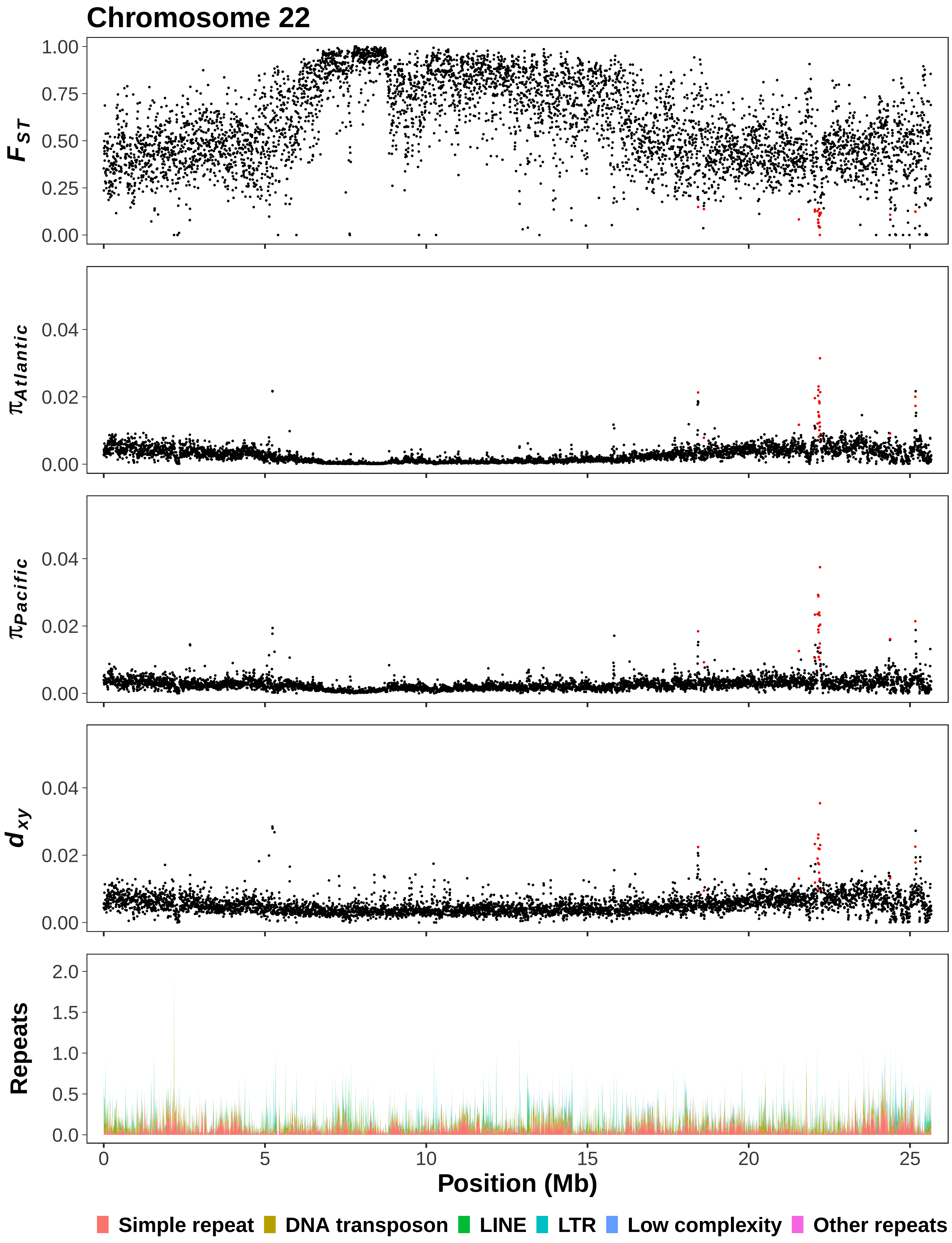

Chromosome 22
$F_{ST}$
1.00
0.75
0.50
0.25
0.00
$\pi_{Atlantic}$
0.04
0.02
0.00
$\pi_{Pacific}$
0.04
0.02
0.00
$d_{xy}$
0.04
0.02
0.00
Repeats
2.0
1.5
1.0
0.5
0.0
0
5
10
15
20
25
Position (Mb)
Simple repeat
DNA transposon
LINE
LTR
Low complexity
Other repeats

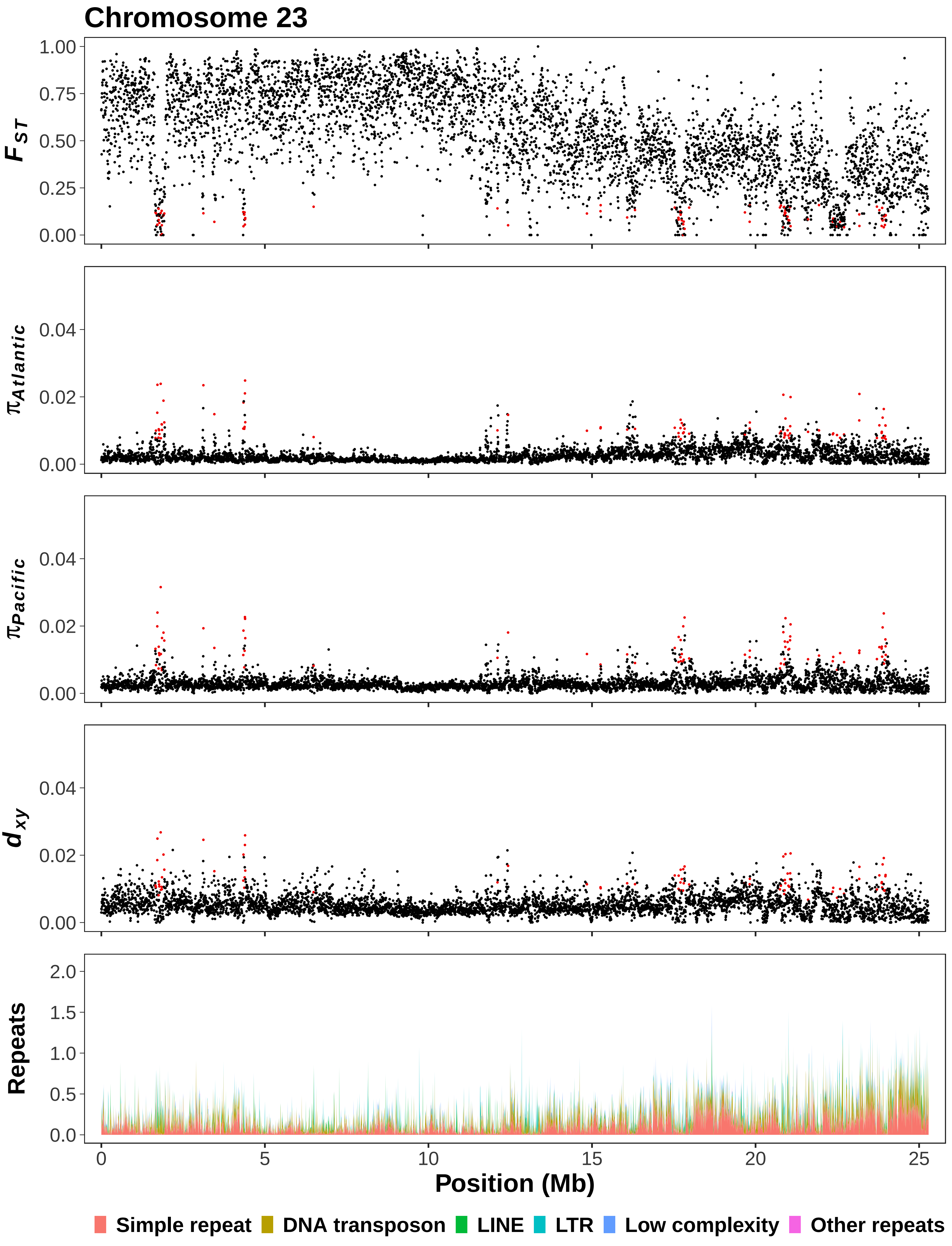

Chromosome 23
$F_{ST}$
1.00
0.75
0.50
0.25
0.00
$\pi_{Atlantic}$
0.04
0.02
0.00
$\pi_{Pacific}$
0.04
0.02
0.00
$d_{xy}$
0.04
0.02
0.00
Repeats
2.0
1.5
1.0
0.5
0.0
0
5
10
15
20
25
Position (Mb)
Simple repeat
DNA transposon
LINE
LTR
Low complexity
Other repeats

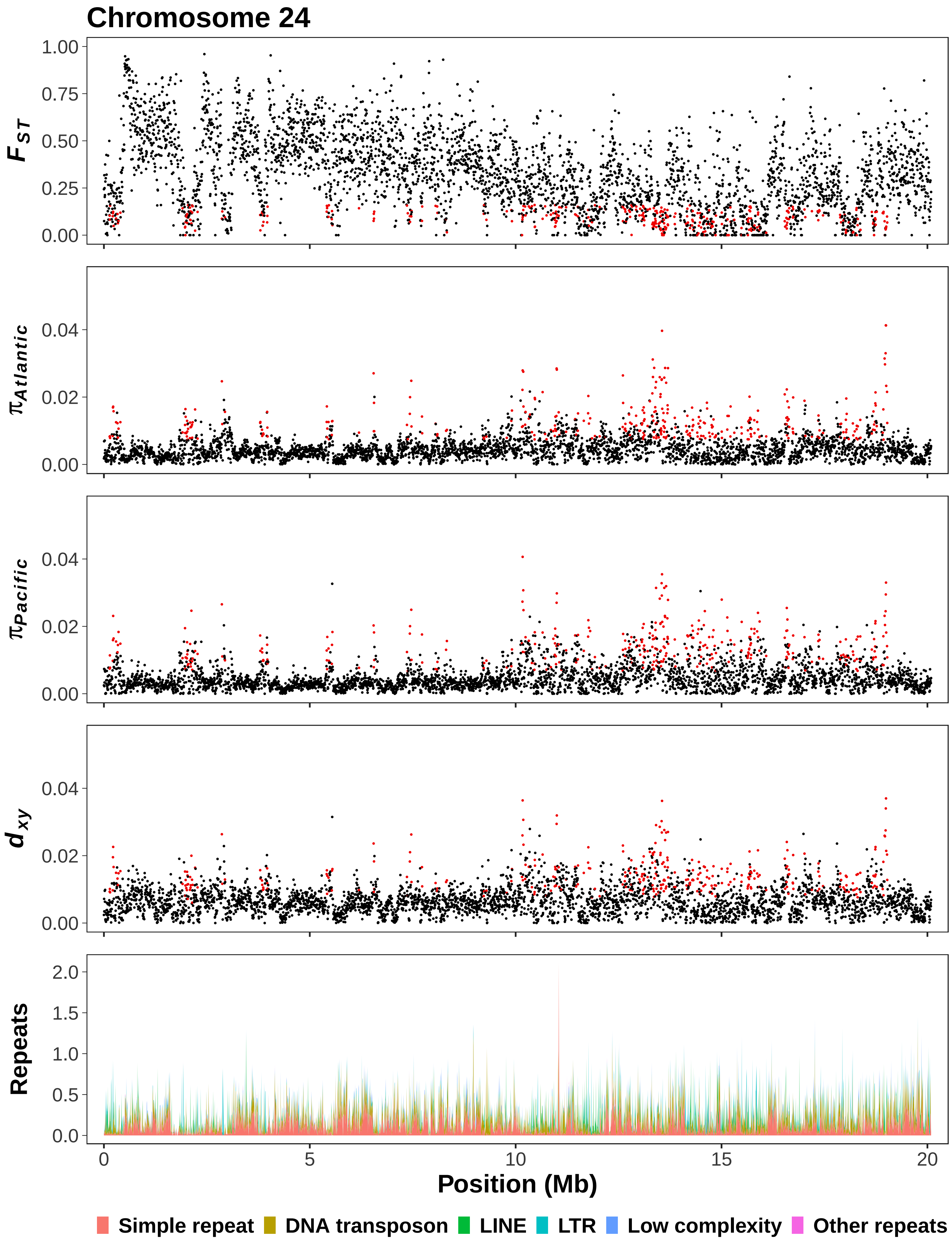

Chromosome 24
$F_{ST}$
1.00
0.75
0.50
0.25
0.00
$\pi_{Atlantic}$
0.04
0.02
0.00
$\pi_{Pacific}$
0.04
0.02
0.00
$d_{xy}$
0.04
0.02
0.00
Repeats
2.0
1.5
1.0
0.5
0.0
0
5
10
15
20
Position (Mb)
Simple repeat
DNA transposon
LINE
LTR
Low complexity
Other repeats

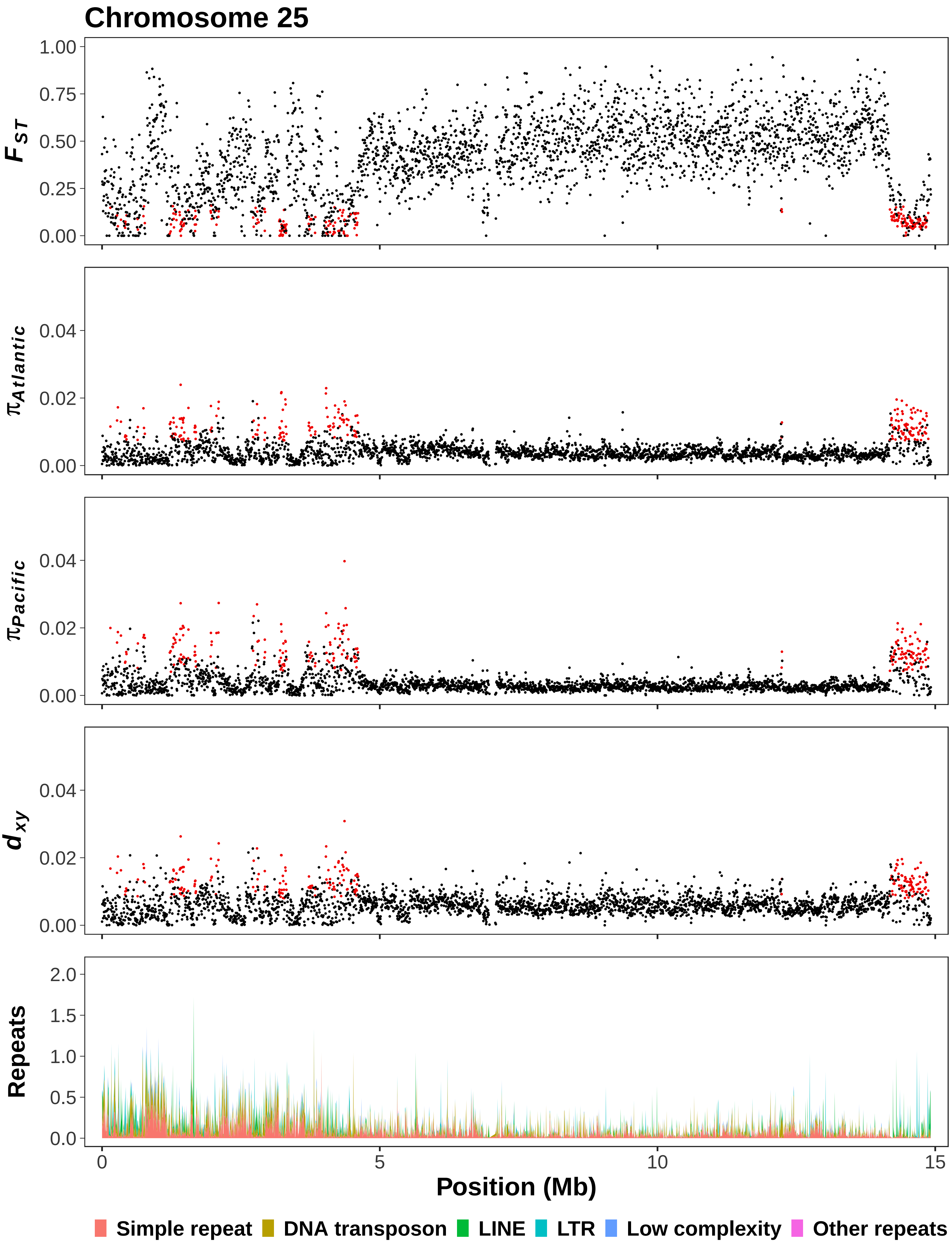

Chromosome 25
$F_{ST}$
1.00
0.75
0.50
0.25
0.00
$\pi_{Atlantic}$
0.04
0.02
0.00
$\pi_{Pacific}$
0.04
0.02
0.00
$d_{xy}$
0.04
0.02
0.00
Repeats
2.0
1.5
1.0
0.5
0.0
0
5
10
15
Position (Mb)
Simple repeat
DNA transposon
LINE
LTR
Low complexity
Other repeats

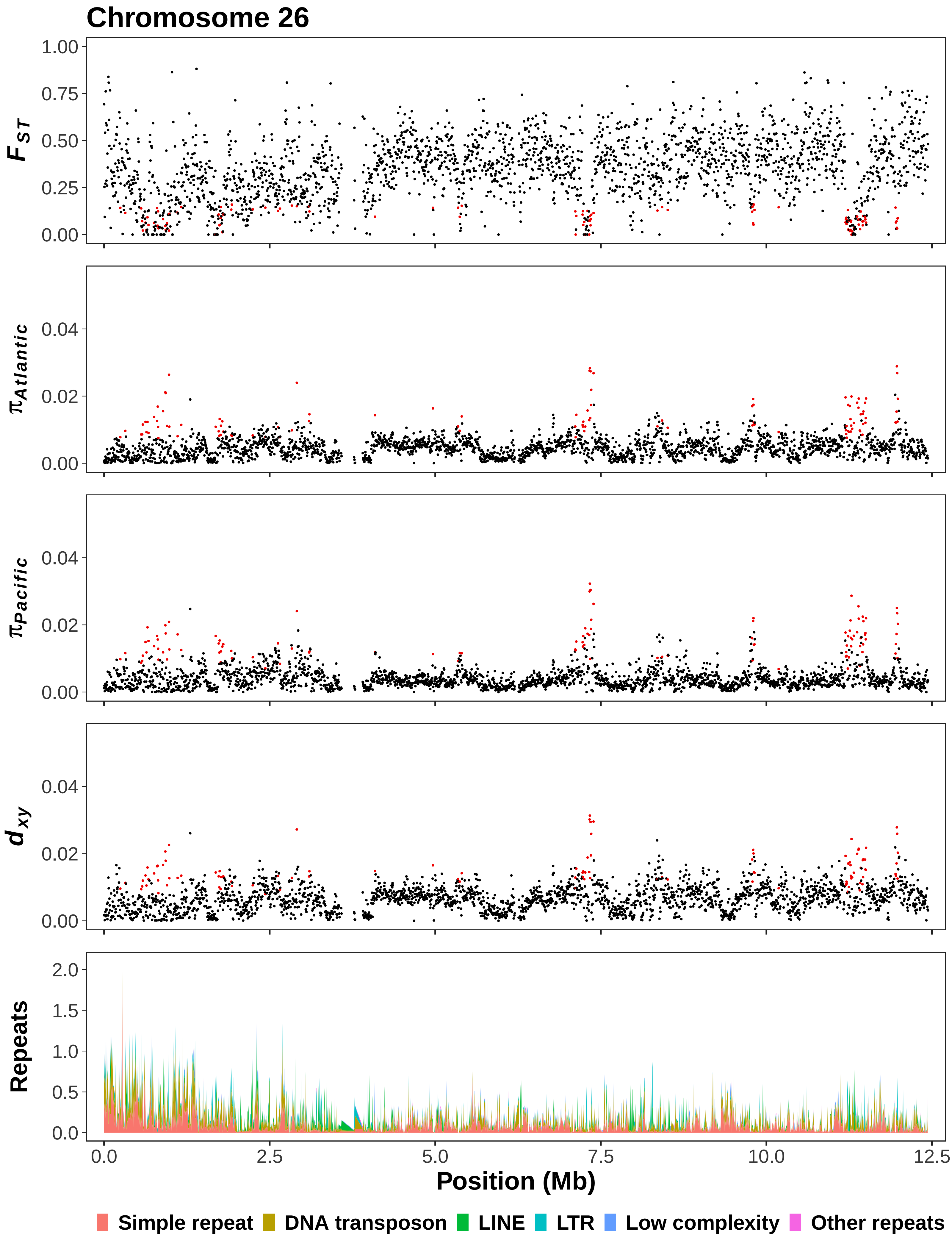
Chromosome 26
$F_{ST}$
$\pi_{Atlantic}$
$\pi_{Pacific}$
$d_{xy}$
Repeats
Position (Mb)
Simple repeat
DNA transposon
LINE
LTR
Low complexity
Other repeats

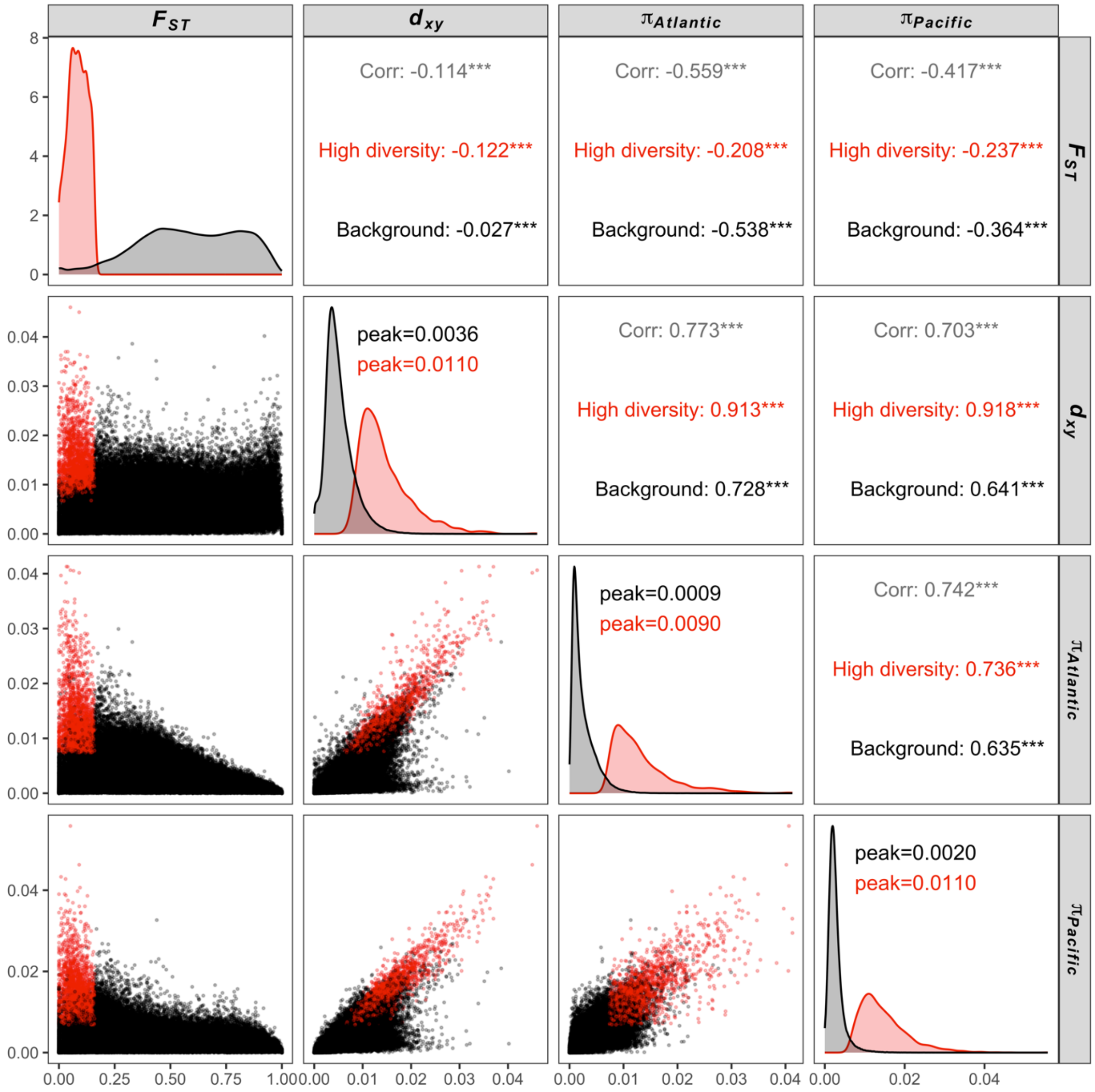


**Figure S2.** Correlation matrix for population genetic diversity parameters between Atlantic and Pacific herring estimated within non-overlapping 5 kb windows. The diagonal shows the density distribution of parameters (the areas under the two density curves in each block are not proportional to the number of bins in each group), the lower triangle displays joint distributions and upper triangle summarises Pearson correlation coefficients between pairs of parameters within high diversity (red) and background (black) windows.

#### A) Biological Process

**1:** homophilic cell adhesion via plasma membrane adhesion molecules
**2:** immune response
**3:** phagocytosis, recognition
**4:** complement activation, classical pathway
**5:** positive regulation of B cell activation
**6:** phagocytosis, engulfment
**7:** regulation of apoptotic process
**8:** B cell receptor signaling pathway
**9:** defense response to bacterium
**10:** innate immune response
**11:** defense response to Gram-negative bacterium
**12:** defense response to Gram-positive bacterium
**13:** negative regulation of alpha-beta T cell proliferation
**14:** proteasomal ubiquitin-independent protein catabolic process

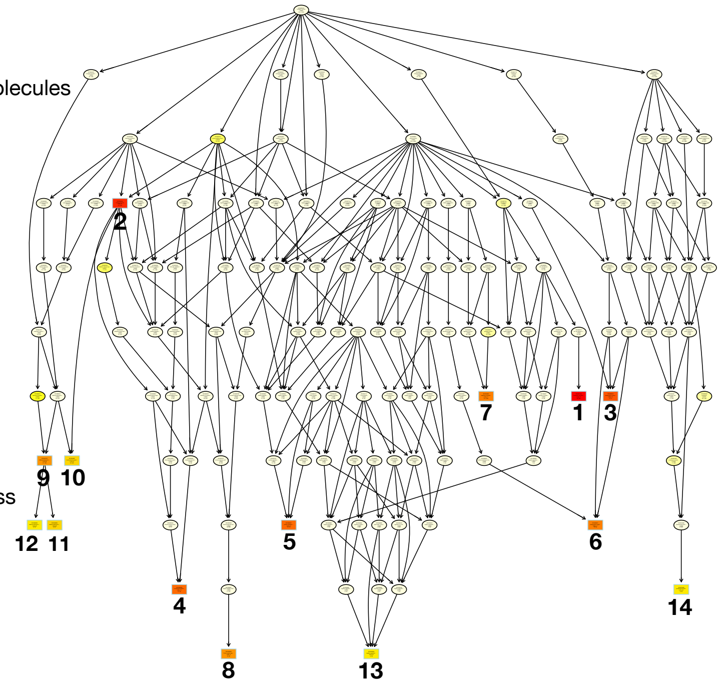


#### B) Molecular Function

**1:** GTP binding
**2:** immunoglobulin receptor binding
**3:** antigen binding
**4:** threonine-type endopeptidase activity
**5:** zinc ion binding

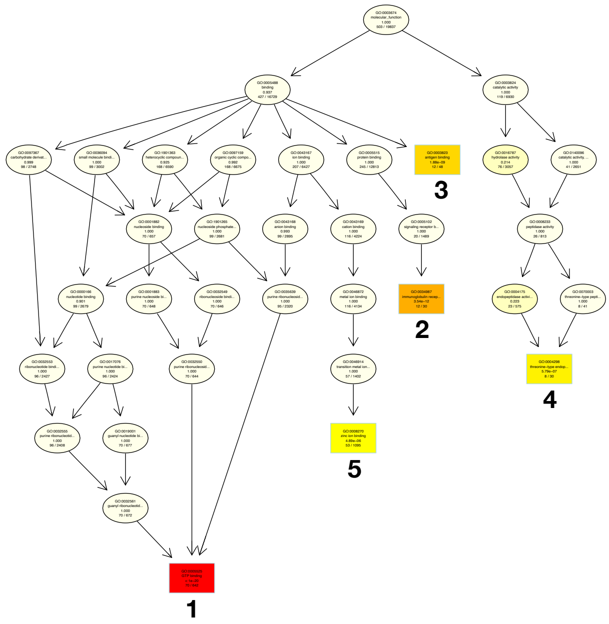


#### C) Cellular component

**1:** intracellular anatomical structure
**2:** MHC class II protein complex
**3:** immunoglobulin complex, circulating
**4:** external side of plasma membrane
**5:** nucleosome
**6:** proteasome core complex

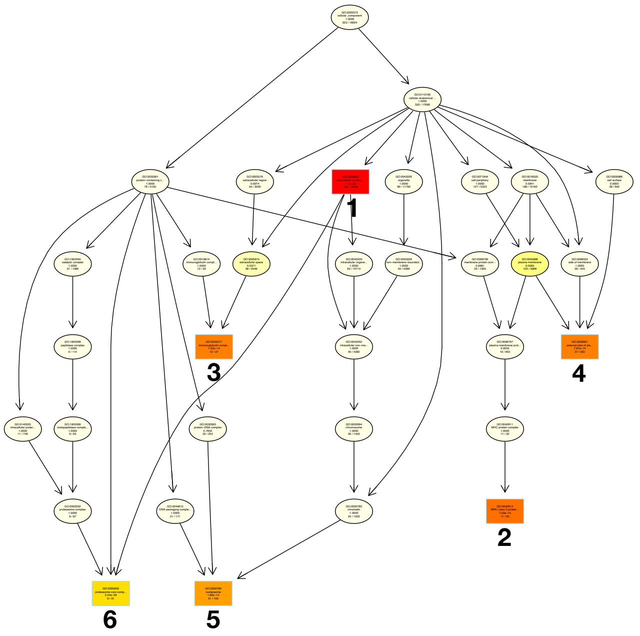

**Figure S3.** Gene ontology subgraphs induced by significant terms in gene set enrichment analysis. Rectangles represent significant biological process (**A**), molecular function (**B**) and cellular component (**C**) terms ($P \leq 0.01$; corrected for multiple testing with the Bonferroni method). The most significant nodes are shown with dark red and light yellow represents the least significant nodes.

(A)

(B)

(C)

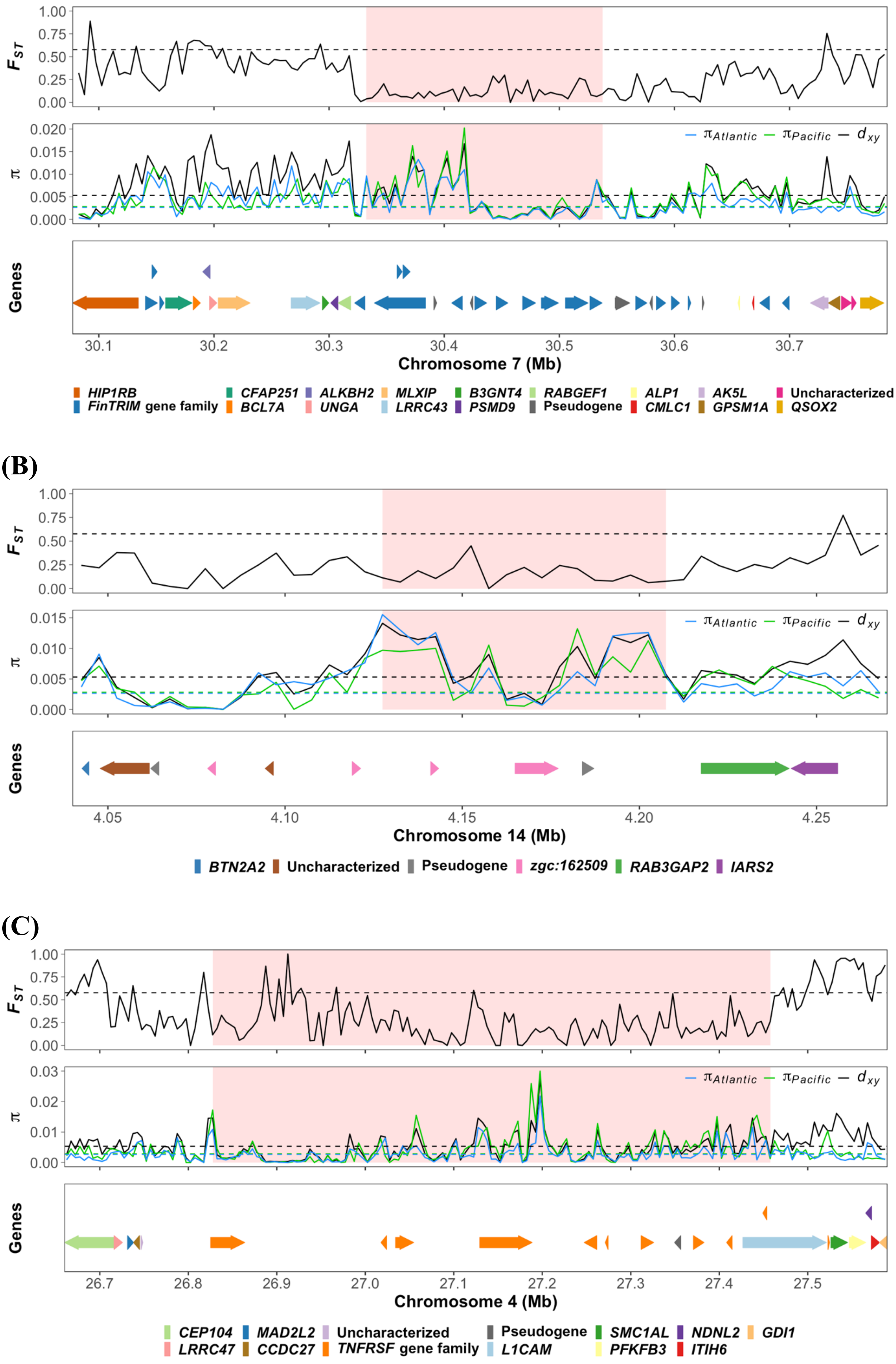

FST
π
πAtlantic
πPacific
dxy
Genes
Chromosome 7 (Mb)
HIP1RB
FinTRIM gene family
CFAP251
BCL7A
ALKBH2
UNGA
MLXIP
LRRC43
B3GNT4
PSMD9
RABGEF1
Pseudogene
ALP1
CMLC1
AK5L
GPSM1A
Uncharacterized
QSOX2
Chromosome 14 (Mb)
BTN2A2
Uncharacterized
Pseudogene
zgc:162509
RAB3GAP2
IARS2
Chromosome 4 (Mb)
CEP104
LRRC47
MAD2L2
CCDC27
Uncharacterized
TNFRSF gene family
Pseudogene
L1CAM
SMC1AL
PFKFB3
NDNL2
ITIH6
GDI1

**Figure S4.** Illustrative examples of immune-related gene family clusters at high diversity regions detected in Atlantic vs. Pacific herring population comparison. The colored area in $F_{ST}$ and $\pi$ plots represents merged high diversity regions in each example. Dashed lines depict respective genome-wide averages. The bottom track in each figure illustrates gene organization in the Atlantic herring reference genome (annotation source: NCBI Clupea harengus Annotation Release 102), indicating a cluster of *FinTRIM* (**A**), *zgc:162509* (**B**) and *TNFRSF* genes (**C**). On the upper row are the nested genes and partially overlapping genes. Color code for genes is given below each figure.

**Table S1.** Samples of Atlantic and Pacific herring used in short-read data analysis.

| Identifier | Location | Latitude | Longitude | Region | Spawning season | Sample size | Salinity (ppt) | Date | Reference |
|---|---|---|---|---|---|---|---|---|---|
| i17 | Bergen | N60°35′ | E05°00′ | Northeast Atlantic Ocean | Spring | 8 | 33 | 20130522 | Martinez Barrio et al. 2016 |
| i34 | Celtic Sea | N51° 59' | W6° 51' | Northeast Atlantic Ocean | Winter | 3 | 35 | 20151201 | Fan et al. 2020 |
| i37 | Downs | N51° 34' | E1° 90' | Northeast Atlantic Ocean | Winter | 3 | 35 | 20161212 | Fan et al. 2020 |
| i10 | Isle of Man | N54° 6' | W4° 37' | Northeast Atlantic Ocean | Autumn | 3 | 35 | 20150930 | Fan et al. 2020 |
| i7 | North Sea | N58°06′ | E6°10′ | Northeast Atlantic Ocean | Autumn | 3 | 35 | 19790805 | Lamichhaney et al. 2017 |
| i31 | Norway | N67° 46' | E9°47' | Northeast Atlantic Ocean | Spring | 3 | 35 | 20170220 | Fan et al. 2020 |
| i1 | Canada(Bonavista Bay) | N48°49′ | W53°20′ | Northwest Atlantic Ocean | Autumn | 2 | 35 | 20140625 | Lamichhaney et al. 2017 |
| i26 | Canada(Fortune Bay) | N47°17′ | W55°38′ | Northwest Atlantic Ocean | Spring | 2 | 35 | 20140526 | Lamichhaney et al. 2017 |
| i5 | Canada(German Banks) | N43°16′ | W66°18′ | Northwest Atlantic Ocean | Autumn | 2 | 35 | 20140828 | Lamichhaney et al. 2017 |
| i30 | Canada(Inner Baie Des Chaleurs) | N48°00′ | W65°51′ | Northwest Atlantic Ocean | Spring | 2 | 35 | 20140508 | Lamichhaney et al. 2017 |
| i3 | Canada(Northumberland Strait) | N45°44′ | W62°36′ | Northwest Atlantic Ocean | Autumn | 2 | 35 | 20140916 | Lamichhaney et al. 2017 |
| i28 | Canada(Northumberland Strait) | N46°19′ | W64°09′ | Northwest Atlantic Ocean | Spring | 2 | 35 | 20140506 | Lamichhaney et al. 2017 |
| i58 | Vancouver, Strait of Georgia | N49°28′ | W123°15′ | Pacific Ocean | Spring | 6 | 35 | 20121124 | Lamichhaney et al. 2017 |

**Table S4.** Examples of gene clusters at regions exhibiting low differentiation and high nucleotide diversity in other fishes. This table highlights gene clusters consisting of a minimum of three genes.

| Species radiation | Approximate coordination | Gene cluster | Annotation source | Reference |
|---|---|---|---|---|
| Midas cichlid | Chr2:19350000-19500000 | interferon-induced very large GTPase 1 | NCBI Archocentrus centrarchus Annotation Release 100 | Kautt et al. 2020 |
| Midas cichlid | Chr2:21600000-22700000 | protocadherin | NCBI Archocentrus centrarchus Annotation Release 100 | Kautt et al. 2020 |
| Midas cichlid | Chr4:17800000-19200000 | Ig heavy chain variable | NCBI Archocentrus centrarchus Annotation Release 100 | Kautt et al. 2020 |
| Midas cichlid | Chr12:5900000-6600000 | GTPase IMAP family member 8-like | NCBI Archocentrus centrarchus Annotation Release 100 | Kautt et al. 2020 |
| Ninespine stickleback | Chr4a:9500000-10500000 | protocadherin | NCBI Pungitius pungitius Annotation Release 100 | Yamasaki et al. 2020 |
| Ninespine stickleback | Chr9:19400000-20400000 | E3 ubiquitin-protein ligase TRIM39<br>carcinoembryonic antigen-related cell adhesion molecule 5 | NCBI Pungitius pungitius Annotation Release 100 | Yamasaki et al. 2020 |
| Ninespine stickleback | Chr13:15800000-16300000 | gastrula zinc finger protein | NCBI Pungitius pungitius Annotation Release 100 | Yamasaki et al. 2020 |
| Ninespine stickleback | Chr14:2200000-2700000 | NACHT, LRR and PYD domains-containing protein 3-like<br>NLR family CARD domain-containing protein 3-like | NCBI Pungitius pungitius Annotation Release 100 | Yamasaki et al. 2020 |

**Table S6**. Nucleotide diversity ($\pi$) based on coding sequences of *CLM2* and *IFIT10* genes in high diversity regions using five assemblies (the reference assembly and four PacBio haplotype assemblies).

| **Genes** | **No. Sequences** | **Size (bp)** | $\pi$ |
|---|---|---|---|
| *CLM2A* | 2 | 468 | 0.004 |
| *CLM2B1* | 4 | 738/741[a] | 0.048 |
| *CLM2B2* | 5 | 690 | 0.032 |
| *CLM2C* | 5 | 696 | 0.016 |
| *IFIT10A* | 5 | 1440 | 0.026 |
| *IFIT10B* | 3 | 1374[b] | 0.001 |

[a]One sequence is 741 bp and three are 738 bp

[b]*IFIT10B* in CS5_h2 is 1370 bp and has a 4 bp deletion compared with other sequences
